# Accurate Genomic Variant Detection in Single Cells with Primary Template-Directed Amplification

**DOI:** 10.1101/2020.11.20.391961

**Authors:** Veronica Gonzalez, Sivaraman Natarajan, Yuntao Xia, David Klein, Robert Carter, Yakun Pang, Bridget Shaner, Kavya Annu, Daniel Putnam, Wenan Chen, Jon Connelly, Shondra Pruett-Miller, Xiang Chen, John Easton, Charles Gawad

## Abstract

Improvements in whole genome amplification (WGA) would enable new types of basic and applied biomedical research, including studies of intratissue genetic diversity that require more accurate single-cell genotyping. Here we present primary template-directed amplification (PTA), a new isothermal WGA method that reproducibly captures >95% of the genomes of single cells in a more uniform and accurate manner than existing approaches, resulting in significantly improved variant calling sensitivity and precision. To illustrate the new types of studies that are enabled by PTA, we developed direct measurement of environmental mutagenicity (DMEM), a new tool for mapping genome-wide interactions of mutagens with single living human cells at base pair resolution. In addition, we utilized PTA for genome-wide off-target indel and structural variant detection in cells that had undergone CRISPR-mediated genome editing, establishing the feasibility for performing single-cell evaluations of biopsies from edited tissues. The improved precision and accuracy of variant detection with PTA overcomes the current limitations of accurate whole genome amplification, which is the major obstacle to studying genetic diversity and evolution at cellular resolution.

## Introduction

Whole genome amplification (WGA) is required for the unbiased sequencing of minute DNA samples. This includes the sequencing of forensic samples(1), ancient genomic fragments(2), unculturable microbes(3), and the genomes of individual eukaryotic cells(4). The sequencing of single human cells has begun providing new insights into the contributions of genetic variation at the cellular level to human health; from establishing roles for somatic mosaicism in human disease(5, 6) to deconvoluting the complexities of cancer clonal evolution(7–9). In addition, these methods are beginning to be applied as diagnostic tools, including detecting disease-initiating genes before the implantation of embryos(10) and the development of higher resolution cancer diagnostics(11). However, single-cell DNA sequencing currently has a limited capacity to detect genetic variation in each cell as a result of the poor data quality produced by existing WGA methods.

Studies aiming to detect genetic variation in single cells have previously been limited to one of two experimental designs. In the first approach, investigators determine copy number variation (CNV) in single cells based on normalized read depth using a WGA method that relies mostly or entirely on PCR amplification of a small portion of the genome, but in a uniform manner(11, 12). Alternatively, investigators identify single nucleotide variation (SNV), but not CNV, using isothermal amplification methods such as multiple displacement amplification (MDA) that amplify most of the genome, but in a highly uneven manner(7, 8, 13). More recently, a method named LIANTI was developed, which utilizes transposases to introduce a T7 promoter for *in vitro* transcription, which is followed by reverse transcription and cDNA amplification(14). LIANTI was shown to have an improved capacity to detect both CNV and SNV in the same cell. However, LIANTI did not overcome some important limitations of previous methods, including a lack of cell-to-cell reproducibility requiring screening of initial amplification products before downstream analyses and limited coverage of both alleles in diploid organisms, resulting in high false negative SNV call rates. In addition, LIANTI has a more complex experimental protocol than previous methods and has not been shown to be amenable to highly parallelized experiments in microfluidic devices.

In the current study, we present a new method we have named Primary Template-Directed Amplification (PTA), which takes advantage of the processivity, strong strand displacement activity, and low-error rate of phi29 polymerase used in MDA. However, in this method, exonuclease-resistant terminators are incorporated into the reaction, creating smaller double-stranded amplification products that undergo limited subsequent amplification. This transforms the reaction from exponential into a quasilinear process with more of the amplification occurring from the primary template (Fig. 1a, Fig. 1b). Moreover, because amplicons are more frequently generated from the original template, errors have limited propagation from daughter amplicons during subsequent amplification, as occurs with MDA (Fig. 1a, left panel). The result is a reaction that, unlike existing WGA protocols, can amplify the genomes of single cells with high coverage breadth and uniformity in a reproducible manner. This produces significantly improved variant calling of all types when compared to all commonly used methods. Moreover, the terminated amplification products can undergo direct ligation of adapters, allowing for the attachment of a cell barcode to the amplification products that can be used for pooling WGA products from all cells for downstream analyses. The result is an easily executed protocol that will enable new applications for studying genetic diversity and evolution at cellular resolution.

**Fig. 1.**
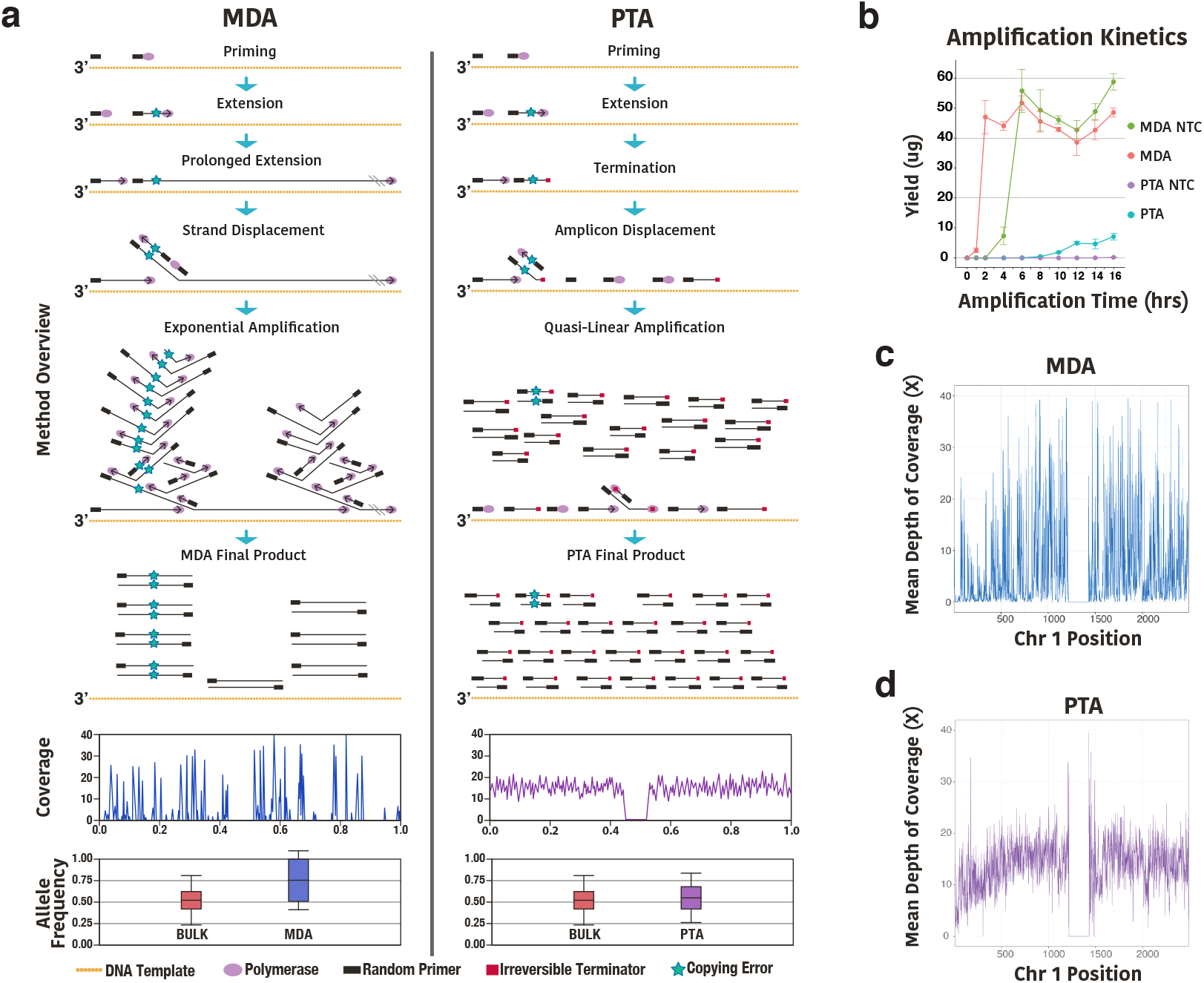
Overview of primary template-directed amplification. **a,** Comparison of MDA to PTA. Both MDA and PTA take advantage of the processivity, strand displacement activity, and low-error rate of the Phi29 polymerase. However, in MDA, exponential amplification at locations where the polymerase first extends the random primers results in overrepresentation of random loci and alleles. In contrast, in PTA, the incorporation of exonuclease-resistant terminators in the reaction result in smaller double-stranded amplification products that undergo limited subsequent amplification, resulting in a quasilinear process with more amplification originating from the primary template. As a result, errors have limited propagation from daughter amplicons during subsequent amplification compared to MDA. In addition, PTA has improved and reproducible genome coverage and uniformity, as well as diminished allelic skewing. **b**, Yield of PTA and MDA reactions over time with either a single cell or no template control (NTC) showing MDA has a much steeper slope as the reaction undergoes exponential amplification. In addition, PTA has very little product detected in NTC samples, compared to similar yields for MDA whether there is a single cell or NTC. **c**, Example of SCMDA coverage and uniformity across chromosome 1 in 100kb bins. **d**, Example of PTA coverage and uniformity across chromosome 1 in 100kb bins. (error bars represent one SD)

## Results

### Irreversible Terminators Produce High-Quality WGA Products

The goal for the method was to improve on the uniformity and coverage of MDA, as it is currently the established method with the lowest error rate and greatest genome coverage breadth. We based our approach on the existing model of MDA where the random overrepresentation of loci and alleles is the result of exponential amplification at locations where the polymerase first extends the random primers. We sought to slow the rate of amplification at those initial sites through the incorporation of dideoxynucleotides to terminate the extension. To accomplish this, we sorted single lymphoblastoid cells from 1000 genomes subject GM12878 whose genome has undergone high-depth whole genome sequencing followed by “platinum” variant calling(15). With the incorporation of standard dideoxynucleotides, we were able to decrease the size of the amplification products (Supplementary Fig. S1a). However, when the amplicons underwent sequencing, we found that the reactions had created poor quality products. The mapping rates and quality scores were 15.0% +/− 2.2 and 0.8 +/− 0.08, respectively (Supplementary Fig. S1b, c). After identifying an overrepresentation of repetitive elements in the data, we hypothesized that amplicons in the genome that were primed first could reprime similar repetitive regions in the genome, as well as similar amplicons in the reaction, creating chimeric sequences that did not align to the human genome.

However, for this model to be accurate, the polymerase would need to remove the terminator from the amplicons prior to extension. To test this model, we incorporated an alpha-thio group into the terminators, which created an exonuclease-resistant phosphorothioate bond when amplification was terminated. Consistent with our model, the irreversible terminators resulted in a significant decrease in the size of the amplification products that were able to undergo direct ligation of adapters that contained a cell barcode (Supplementary Fig. S1a). More importantly, there was a vast improvement in the quality of the amplification products (Supplementary Fig. S1b, c). For example, compared to the reversible terminators, the percent of reads mapped increased from 15.0+/−2% to 97.9+/−0.62% and the mapping quality scores increased from 0.8+/−0.08 to 46.3+/−3.18 (Supplementary Fig. S1b, c). Also of importance, no template control reactions with the irreversible terminators did not have detectable amplification products, showing that the nonspecific amplification that occurs with MDA is suppressed (Fig. 1b). Together, these modifications resulted in a dramatic change in the characteristics of the reaction, resulting in the creation of our new WGA method which we have named PTA.

### PTA Has Increased and Reproducible Genome Coverage Breadth and Uniformity

We then performed comprehensive comparisons of PTA to common single-cell WGA methods. To accomplish this, we performed PTA and an improved version of MDA called single-cell MDA(16) (SCMDA) on 10 GM12878 cells each. In addition, we compared those results to cells that had undergone amplification with DOP-PCR(17), General Electric MDA(18), Qiagen MDA, MALBAC(19), LIANTI(14), or PicoPlex(20) using data that were produced as part of the LIANTI study.

To normalize across samples, raw data from all samples were aligned and underwent pre-processing for variant calling using the same pipeline. The bam files were then randomly subsampled to 300 million reads each prior to performing comparisons. Importantly, the PTA and SCMDA products were not screened prior to performing further analyses while all other methods underwent screening for genome coverage and uniformity before selecting the highest quality cells that were used in subsequent analyses. Of note, SCMDA and PTA were compared to bulk diploid GM12878 samples while all other methods were compared to bulk BJ1 diploid fibroblasts that had been used in the LIANTI study. As seen in Supplementary Fig. S2, PTA had the highest percent of reads aligned to the genome, as well as the highest mapping quality. PTA, LIANTI, and SCMDA had similar GC content, all of which were lower than the other methods. PCR duplication rates were similar across all methods (Supplementary Fig. S2d). Interestingly, by relying more on the primary template for amplification, PTA enabled smaller circular templates, such as the mitochondrial genome, to compete with the larger nuclear chromosomes to undergo greater relative amplification when compared to all other WGA methods (Supplementary Fig. S3).

We then sought to compare the coverage breadth and uniformity of all methods. Examples of coverage plots across chromosome 1 are shown for SCMDA and PTA, where PTA is shown to have significantly improved uniformity of coverage compared to SCMDA (Fig. 1c, d). Coverage rates were then calculated for all methods using increasing number of reads. PTA approaches the two bulk samples at every depth, which is a significant improvement over all other methods (Fig. 2a). We then used two strategies to measure coverage uniformity. The first approach was to calculate the coefficient of variation of coverage at increasing sequencing depth where PTA was found to be more uniform than all other methods (Fig. 2b). The second strategy was to compute Lorenz curves for each subsampled bam file where PTA was again found to have the greatest uniformity (Fig. 2c). To measure the reproducibility of amplification uniformity, we calculated Gini Indices to estimate the difference of each amplification reaction from perfect uniformity(21). PTA was again shown to be reproducibly more uniform than the other methods (Fig. 2d).

**Fig. 2.**
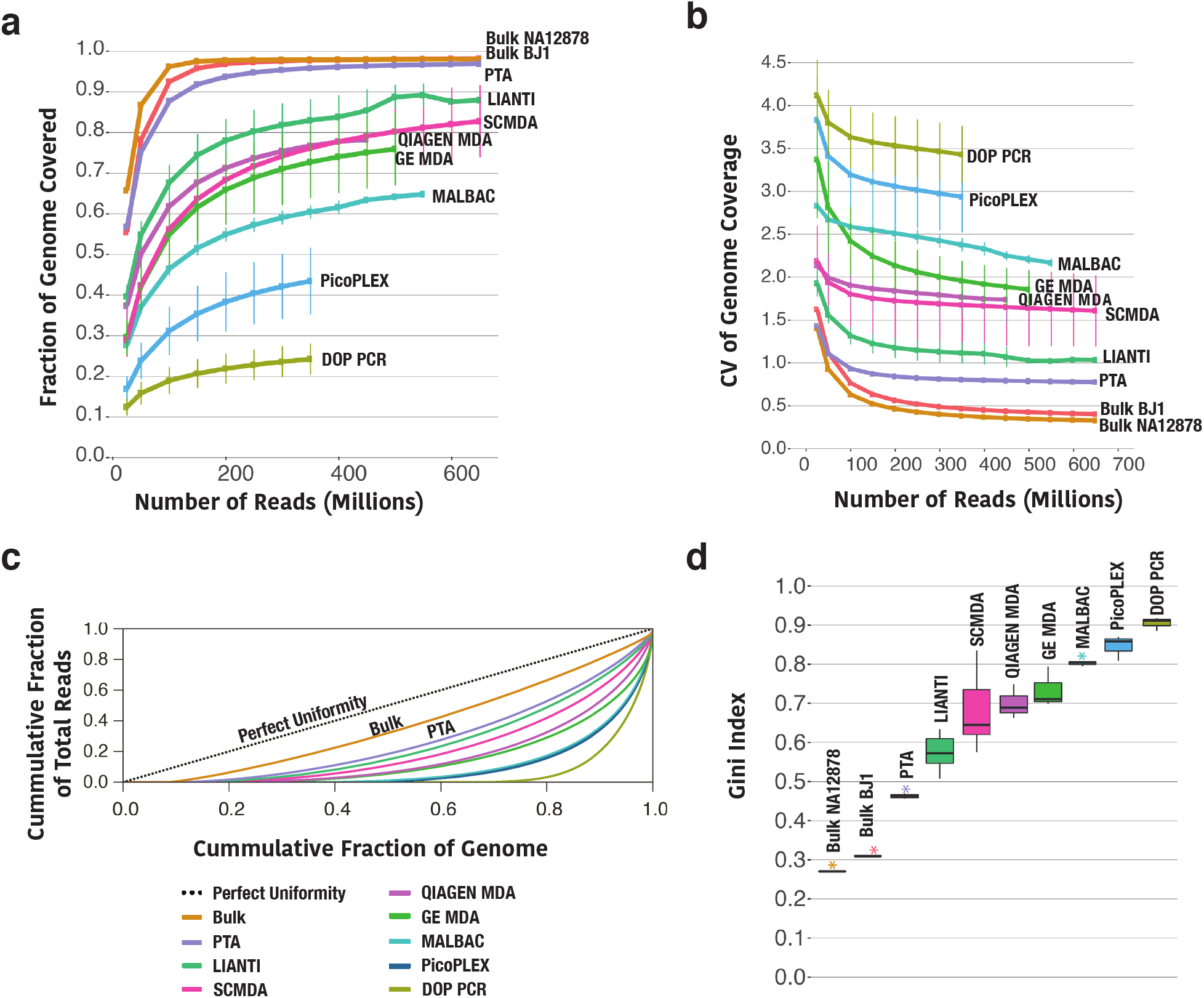
Single cell genome coverage breadth and uniformity of different WGA methods. PTA and SCMDA were performed on random GM12878 cells (n=10, dot represents the mean of 10 individual cells) while DOP-PCR (n=3), GE MDA (n=3), Qiagen MDA (n=3), MALBAC (n=3), LIANTI (n=11), or PicoPlex (n=3) were performed on selected BJ1 cells as part of the LIANTI study. **a**, Genome coverage comparison across different methods at increasing number of single end sequencing reads. PTA approaches the genome coverage obtained in both bulk samples at every sequencing depth. Note that 600 million 150bp single reads represents about 30X whole genome coverage.**b**, Genome coverage uniformity as measured by the coefficient of variation at increasing sequencing depth. **c**, Genome coverage uniformity as measured by the Lorenz curves. The diagonal line represents perfectly uniform genome coverage. The further a curve deviates from the diagonal line, the more bias in genome coverage. PTA is the WGA method that most closely approximates to the genome coverage uniformity obtained from bulk sequencing. The bulk curve was calculated from reads of unamplified bulk GM12878 sample. **d**, Reproducibility of amplification uniformity. The Gini Index measures the departure from perfect uniformity. Smaller standard deviation and lower Gini Index values were measured in PTA samples (purple asterisk), as compared to the other WGA methods. (for dot and line plots error bars represent one SD, for boxplots center line is the median; box limits represent upper and lower quartiles; whiskers represent 1.5x interquartile range; points show outliers)

### PTA Has Significantly Improved SNV Calling Sensitivity

To determine the effects of these differences in the performance of the amplification on SNV calling, we compared the variant call rates for each of the methods to the corresponding bulk samples at increasing sequencing depth. To estimate sensitivity, we calculated the percent of variants called in corresponding bulk samples that had been subsampled to 650 million single reads that were found in each cell at a given sequencing depth (Fig. 3a). The improved coverage and uniformity of PTA resulted in the detection of greater than 90% of variants compared 65-70% of variants detected with LIANTI, which was the next most sensitive method. To measure the difference in amplification between alleles, we then calculated the allele frequencies of variants that had been called heterozygous in the bulk sample where we found that PTA had significantly diminished allelic dropout and skewing at those heterozygous sites (Fig. 3d). This finding supports the assertion that PTA not only has more even amplification across the genome, but also more evenly amplifies both alleles in the same cell. Interestingly, LIANTI shows similar allelic skewing rates to MDA even though it was shown to amplify across the genome in a more even manner.

**Fig. 3.**
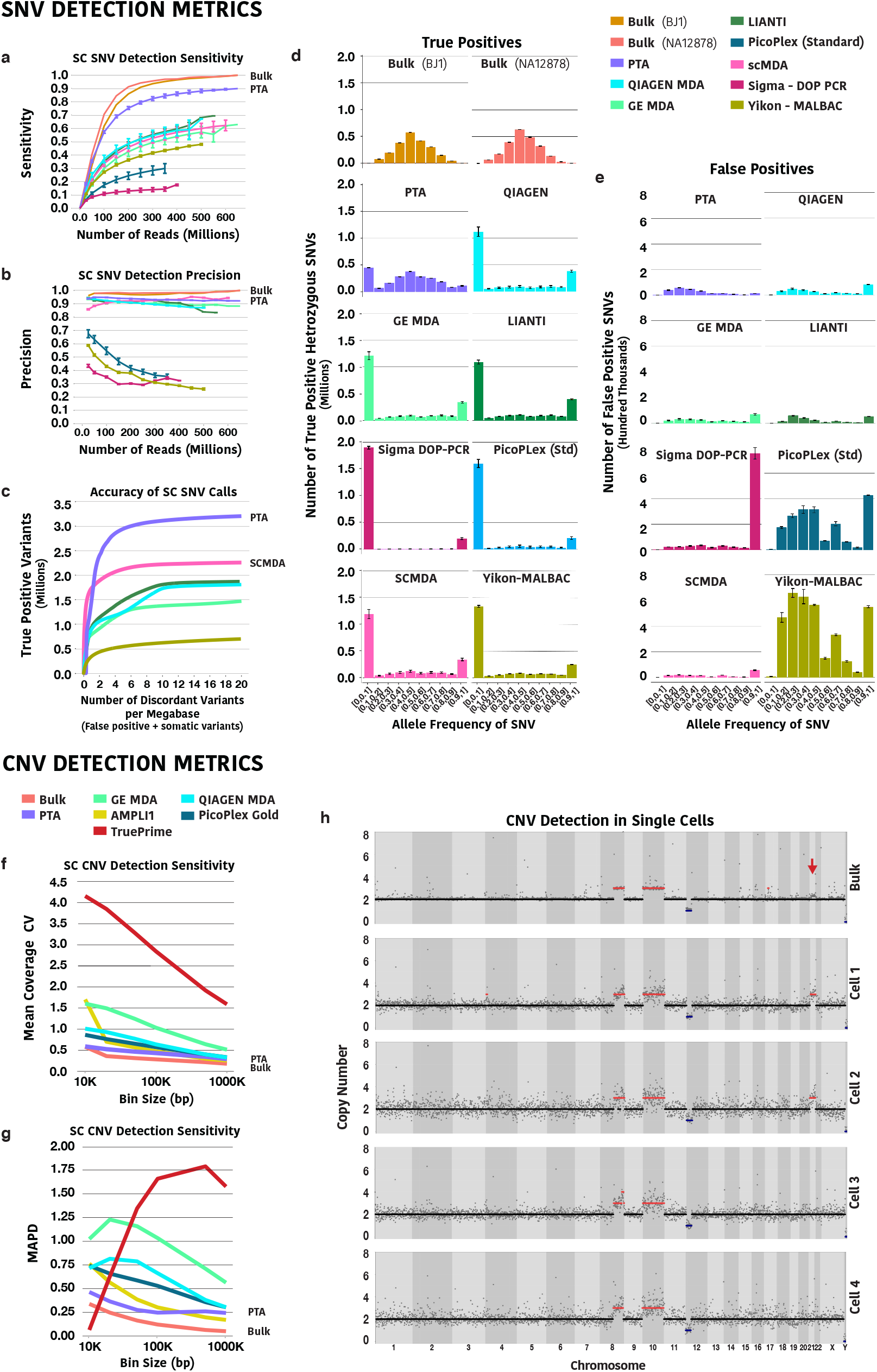
Improved PTA coverage metrics increases variant detection accuracy. a, Comparison of the SNV detection sensitivity in single cells. The SNV sensitivity of each method was calculated as the ratio of the variants identified in each cell by that method to the variants identified in the corresponding unamplified bulk sample at a given sequencing depth. The increased genome coverage, as well as the more uniform distribution of the reads across the genome significantly improves the detection of SNVs by PTA over all other single cell WGA methods. **b**, Comparison of SNV calling precision in single cells. Discordant calls in single cells (FP plus somatic variants) were defined as variant calls in single cells not found in the corresponding bulk samples. Methods using low temperature lysis and/or isothermal polymerase produced significantly lower discordant calls than methods using thermostable polymerases. **c,** Summary of SNV calling accuracy for each method at decreasing VQSLOD quality score from GATK. **d,** Comparison of allele dropout and frequencies of SNVs called heterozygous in the bulk sample. PTA more evenly amplifies both alleles in the same cell, resulting in significantly diminished allelic dropout and skewing. **e**, Comparison of discordant variant call allele frequencies. The quasilinear amplification and suppression of error propagation in PTA result in lower discordant calls than all other methods. **f,** Mean coefficient of variation (CV) of coverage at increasing bin size in a primary leukemia sample using the latest commercially available kits as an estimate of CNV calling accuracy (corresponding coverage and SNV calling metrics for these methods in these samples presented in Fig. S5, n=5 for each method) **g,** Mean median absolute pairwise distance (MAPD) at increasing bin size as a second estimate of CNV calling sensitivity (n=5 for each method). **h,** Example of CNV profiles of PTA product from single cells and DNA from the corresponding bulk sample. The red arrow represents an area where subclonal gain of chromosome 21 was suggested but not called in the bulk sample, while three of eight cells were found to have the same alteration (additional cells and samples are presented in Figs. S7 and S8). (for dot and line plots error bars represent one SD, for boxplots center line is the median; box limits represent upper and lower quartiles; whiskers represent 1.5x interquartile range; points show outliers).

To estimate the precision of SNV calls, the variants called in each single cell not found in the corresponding bulk sample were considered false positives. As previously reported, the lower temperature lysis of SCMDA significantly reduced the number of false positive variant calls (Figs. 3b, 3e)(16). Interestingly, the methods that use thermostable polymerases (MALBAC, PicoPlex, DOP-PCR) showed further decreases in the SNV calling precision with increasing sequencing depth, which is likely the result of the significantly increased error rate of those polymerases compared to phi29 DNA polymerase(22). In addition, the base change patterns seen in the false positive calls also appear to be polymerase-dependent (Supplementary Fig. S4). Further, for PTA, true and false positive calls did not appear to be related to read depth at a given allele frequency as is typically seen for germline variants called from bulk samples, suggesting deeper sequencing would not capture additional variants (Supplementary Fig. S5). As seen in Fig. 3b, our model of suppressed error propagation with PTA is supported higher precision of SNV calling with PTA compared to standard MDA protocols. To examine the tradeoff in sensitivity and precision for a given quality metric, we examined SNV calling accuracy with GATK4, Monovar(23), and SCCaller(16) where we again found PTA to have significantly improved SNV calling sensitivity over all other methods while retaining high precision (Figs. 3c, S7). With these analyses, we also noted that GATK4 and Monovar performed similarly for PTA data while SCCaller had the lowest sensitivity and precision. We then examined the allele frequencies of the discordant variant calls where we see that PTA has the lowest allele frequencies, which is again consistent with our model of suppressed error propagation with PTA (Fig. 3d).

### Calling SNV and CNV with PTA in Primary Cancer Cells

To insure our findings were not due to artifacts produced from comparing data from two different studies while also adding evaluations of the latest commercially available WGA methods, we then used primary leukemia cells to perform further validation studies of PTA for SNV and copy number variation (CNV) calling. This version of the PTA protocol incorporated a low temperature lysis step to suppress higher temperature-medaited cytosine deamination and showed similar genome coverage breadth and uniformity (Supplementary Fig. S6). PTA also remained the most sensitive method for SNV calling at all sequencing depths, as well as the highest SNV calling precision by changing to the low temperature lysis. Of note, the latest version of the REPLI-g method had improved coverage breadth, uniformity, and reproducibility, resulting in improved variant calling sensitivity. The methods that rely on PCR (Ampli1, PicoPlex Gold) also continued to show decreased precision at increasing sequencing depths, although the drop in precision was significantly improved over MALBAC and the previous version of PicoPlex.

To estimate the accuracy of calling CNV of different sizes for each method, we subsampled each bam file to 300 million single reads and measured the CV, as well as the median absolute pairwise distance (MAPD), at increasing bin sizes (Figs. 3f, 3g). We found that PTA again had the lowest CV and MAPD compared to all other WGA methods at every bin, with the exception of the MAPD of Ampli1 with large bin sizes (Fig. 3g). Notably, TruePrime had a lower caculated MAPD at smaller bin sizes due to the high number of bins with zero coverage.

This particular leukemia sample had known CNV on chromosomes 8, 10, and 12. CNV analysis found the two gains and single deletion in each cell. Interestingly, the bulk data suggested there may be a gain of chromosome 21 that was not called in the bulk sample (Fig. 3h). Three of the eight single cells were indeed found to have gain of chromosome 21, suggesting single-cell CNV profiling may be more sensitive, as well as better a better strategy for estimating the percent of cells in a tissue that have a given copy number change (Figs. 3h, S8). Profiling of cells from four additional samples provide further evidence that PTA can be used to reproducibly call CNV in different contexts (Fig. S9).

### Accurately Measuring SNV Rates in Kindred Cells

One important limitation of our previous strategy using discordant variant calls from the bulk sample to measure variant calling precision was that we could not differentiate false positive calls from true somatic variants. To provide even more accurate measurements of variant calling sensitivity and precision, we performed kindred cell studies by plating single CD34^+^ CB cells into a single well, followed by expansion for five days (Fig. 4a). Single cells were then reisolated from that culture to compare variant calling of cells that were almost genetically identical. Further, using the bulk as a reference, we were able to discriminate between germline, false positive, and somatic variant calls (Fig. 4b). With this approach, and again using the bulk sample as the ground truth, we determined our variant calling precision with the low temperature protocol increased to 99.9% (Fig. 4c). Further, most of these primary cells had similar or improved variant detection sensitivity. However, there was one notable cell that had significantly lower variant calling sensitivity, which could be the result of manually manipulating fragile primary cells. Interestingly, the false positive variants in the highest quality cells had a skewing to lower allele frequencies, which could be explained by these rapidly dividing cells being tetraploid in late S or G2/M phase of the cell cycle with only one of four alleles acquiring a copying error (Supplementary Fig. S10). Interestingly, the homozygous false positive calls cluster at specific locations while the heterozygous calls do not. This could be the result of loss or lack of template denaturating at one allele at those locations during the amplification, which does not appear to be dependent on the GC content of the genomic region, although those locations are clustered near the centromere or telomere (Supplementary Fig. S11).

**Fig. 4.**
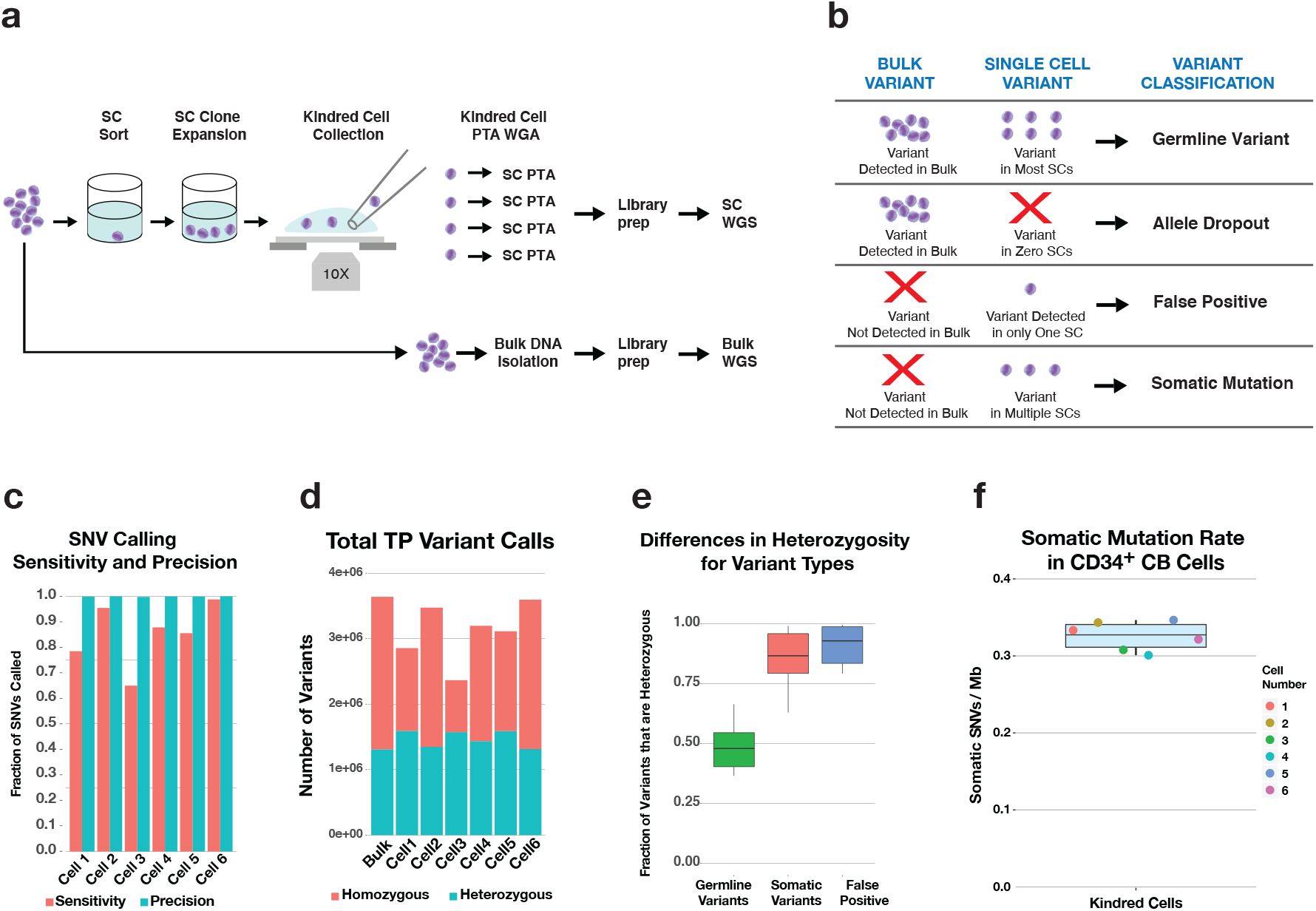
Using kindred cells to accurately measure SNV types. **a**, Overview of strategy for kindred cell experiment where single cells are plated and cultured prior to reisolation, PTA, and sequencing of individual cells. **b**, Strategy for classifying variant types by comparing bulk and single cell data. **c**, SNV calling sensitivity and precision for each cell using the bulk as the gold standard. **d,** Total number of true positive variants detected in each cell. **e**, Percent of variants that were called heterozygous for different variant classes. **f**, Measured somatic variant rates in a single CD34^+^ human cord blood cell after focusing on the highest quality variant calls (DP>=10, GQ>=20, allele frequency>=0.35) of 0.33 +/− 0.02 somatic SNVs per CD34+ cord blood cell.

We then determined that most false positive and somatic variants were called heterozygous, which is consistent with the model that only one allele is mutated as a result of copying errors or during development, respectively (Fig. 4e). Finally, we focused on the highest quality variant calls (DP>=10, GQ>=20, allele frequency>=0.35) and used the estimated genomewide SNV calling sensitivity of those variants to reproducibly estimate a per cell somatic SNV rate of 0.33 +/− 0.02 per Mb, or about 1000 somatic SNVs per hematopoietic stem cell genome (Fig. 4f). Taken together, these data confirm that PTA can accurately detect a much larger number of variants across the genome of single cells than existing WGA methods with high precision, and that the highest quality variants can be used to accurately measure per cell mutation rates.

### Creating Single Cell Maps of Mutagen-Genome Interactions at Base Pair Resolution

To present an example of the new types of studies that can be performed with PTA, we developed a mutagenicity assay that provides a framework for performing high-resolution, genome-wide human toxicogenomics studies. The Ames test revolutionized our capacity to measure the mutagenicity of environmental compounds(24), and is still widely used to evaluate the mutagenicity of industrial compounds, agrochemicals, and pharmaceuticals. However, the system relies on the indirect measurement of DNA damage in which histidine-dependent bacteria need to acquire revertant mutations that allow them to survive in a histidine-deficient environment. Still, these measurements only provide a rough estimate of the mutation numbers and patterns in bacteria. Single-cell mutagenesis assays have been developed, but provide an imcomplete picture as a result of limited SNV detection sensitivity in cells after MDA(25). To provide more comprehensive high-resolution details of the mutagenicity of compounds in human cells, we developed a system we have named direct measurement of environmental mutagenicity (DMEM) where we expose human cells to an environmental compound, perform high-quality single cell variant calling with PTA, and create a map of mutagen-genome interactions that occurred while each cell was alive.

For our proof-of-concept study, we exposed human umbilical CB cells that express the stem/progenitor marker CD34 to increasing concentrations of the direct mutagen N-ethyl-N-nitrosourea (ENU) or D-mannitol (MAN), a compound commonly used at toxic doses as a negative control in mutagenesis studies. ENU is known to have a relatively low Swain-Scott substrate constant and has consequently been shown to predominantly act through a two-step SN1 mechanism that results in preferential alkylation of O4-thymine, O2-thymine, and O2-cytosine(26). Through limited sequencing of target genes, ENU has also been shown to have a base change preference for T to A (A to T), T to C (A to G), and C to T (G to A) changes in mice (27), which significantly differs from the mutation pattern seen in *E. coli* (28).

Consistent with these studies, we measured a dose-dependent increase in mutation numbers in each cell, where a similar number of mutations were detected in the lowest dose of ENU compared to either vehicle control or toxic doses of mannitol (Fig. 5a). Also consistent with previous work in mice, we see the most common mutations are T to A (A to T), T to C (A to G), and C to T (G to A). Interestingly, we do detect the other three types of base changes, although C to G (G to C) transversions appear to be rare (Fig. 5b). We hypothesized that 5-methylcytosine does not undergo alkylation by ENU due to inaccessibility in heterochromatin or as a result of unfavorable reaction conditions with 5-methylcytosine compared to cytosine. To test the former hypothesis, we compared the locations of the mutation sites to known DNase I hypersensitive sites in CD34^+^ cells that were catalogued by the Roadmap Epigenomics Project (29). As seen in Fig. S12, we do not see an enrichment of cytosine variants in DNase I hypersensitivity sites. To further support our model of ENU mutagenesis being independent of genome structure, we also found that genomic feature annotation for the variants is not significantly different from the annotation of those features in the genome (Fig. S13).

**Fig. 5.**
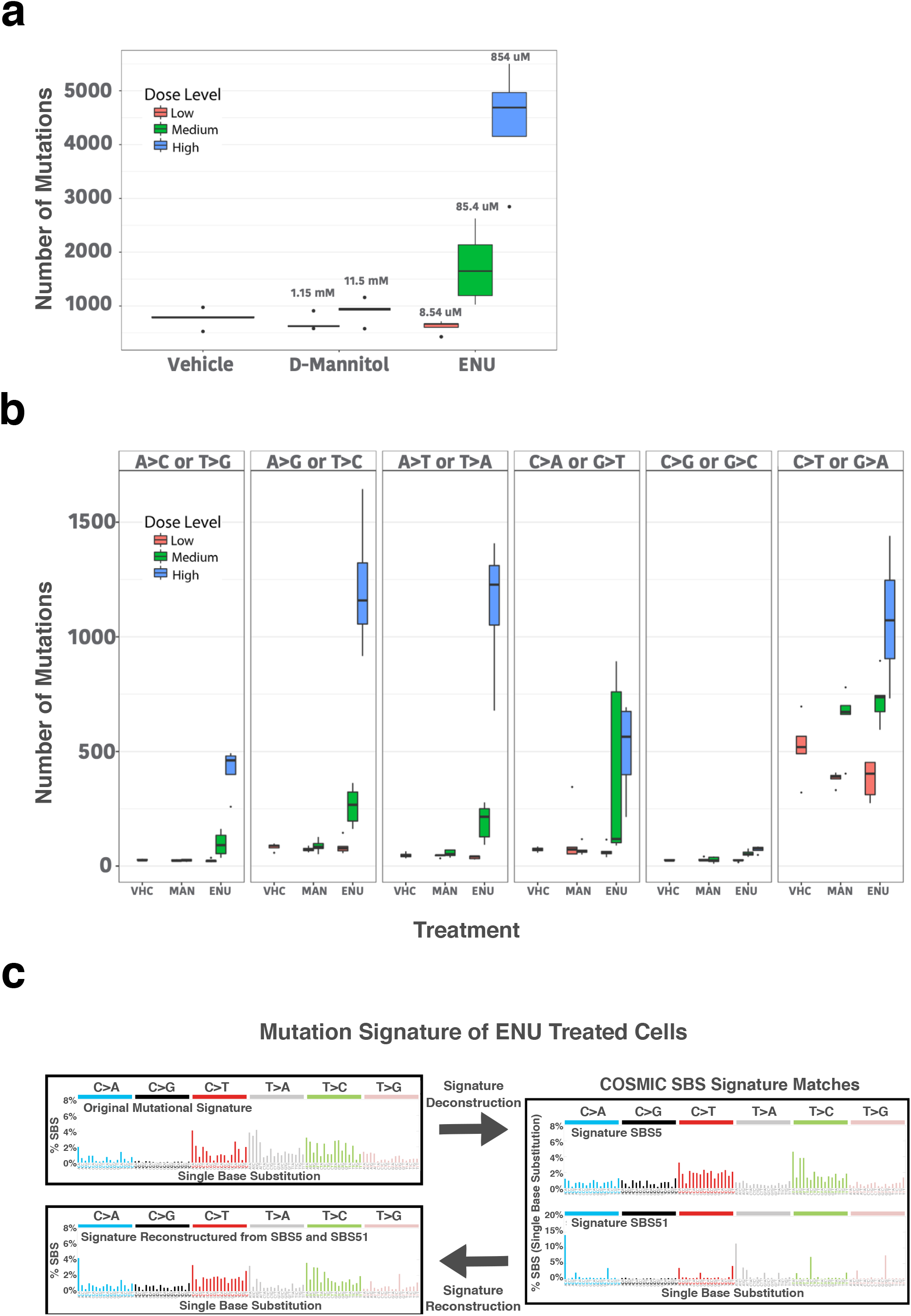
Measuring mutagenicity *In vivo* at single-cell resolution using direct measurement of environmental mutagenicity (DMEM). In the DMEM assay, human cells are exposed to a test compound. Exposed cells then undergo PTA and single-cell sequencing to create a map of genome-mutagen interactions in living cells. a, Single cells exposed to N-ethyl-N-nitrosourea (ENU) or D-mannitol (MAN) show a dose-dependent increase in ENU-induced mutagenesis. (n=5 for all samples except highest dose of ENU where n=4). b, Base change preference by ENU and D-mannitol identify the previously recognized T to C (A to G) and T to A (A to T) base changes as being most common with ENU exposure. c, Identification of a unique ENU mutational signature which was deconvoluted into known COSMIC single base substitution signatures. The relative proportion of COSMIC signatures were then used to reconstruct a signature that can then be compared to the original ENU signature to identify which changes are captured by the approach. (for boxplots center line is the median; box limits represent upper and lower quartiles; whiskers represent 1.5x interquartile range; points show outliers)

We then created a signature of ENU-induced mutagenesis using SigProfiler(30), which tries to deconvolute a given signature into known COSMIC signatures. The single bases substitution signature was most similar to COSMIC signatures SBS5 and SBS51 (Fig. 5c). However, when those signatures were combined and compared to the original ENU signature, there are based changes that are clearly different and not captured by the COSMIC signatures, including an overrepresentation of cytosine to guanine changes and an underrepresentation of thymine to adenine mutations. A similar strategy was used to create a double base substitution signature (Supplementary Fig. S14).

### Measuring Rates and Locations of CRISPR Off-Target Activity in Single Human Cells

The continued development of genome editing tools shows great promise for improving human health, from correcting genes that result in or contribute to the formation of disease(31) to the eradication of infectious diseases that are currently incurable(32, 33). However, the safety of these interventions remain unclear as a result of our incomplete understanding of how these tools interact with and permanently alter other locations in the genomes of the edited cells. Methods have been developed to estimate the off-target rates of genome editing strategies, but most of the tools that have been developed to date interrogate groups of cells together, limiting the capacity to measure the per cell off-target rates and variance between cells, as well as to detect rare editing events that occur in a small number of cells(34–36). Single cell cloning of edited cells has been performed, but could select against cells that acquire lethal off-target editing events and is impractical for many types of primary cells (37).

Taking advantage of the improved variant calling sensitivity and specificity of PTA, here we present a strategy for making quantitative measurements of CRISPR-mediated genome editing with specific guide RNAs (gRNA) in single cells. We utilized three cell types for these studies: U20S osteosarcoma cell line, primary hematopoietic CD34+ CB cells, and embryonic stem (ES) cells. In addition, we employed two previously described gRNAs, one that is known to be precise (EMX1), and one that is known to have high levels of off-target activity (VEGFA)(35). To identify indels with high specificity, we restricted our variant calling to genome locations that had a perfect match to the PAM sequence and up to five mismatches to the protospacer (Fig. 6a).

**Fig. 6.**
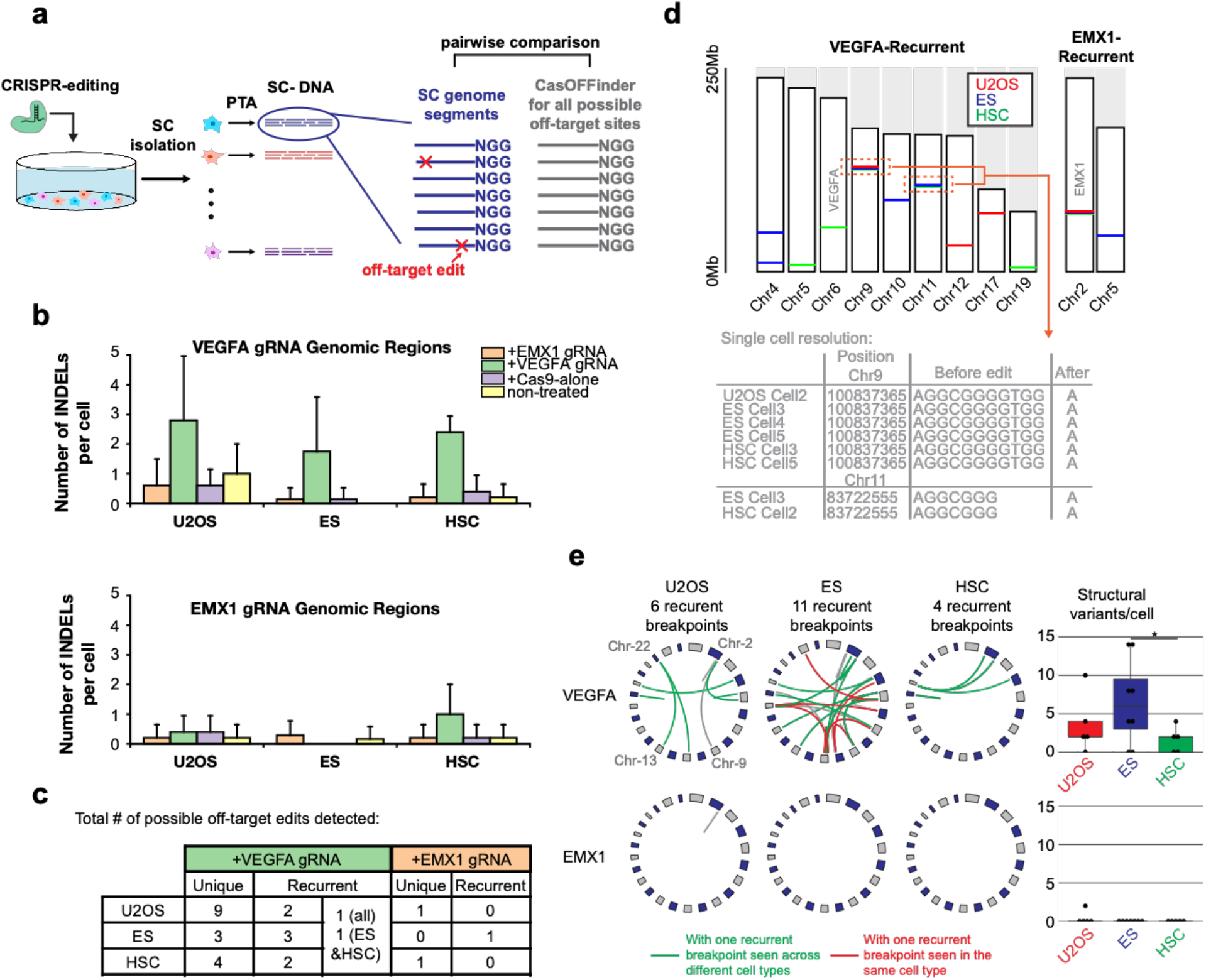
Measuring off-target activity of genome editing strategies at single-cell resolution. **a**, Overview of experimental and computational strategy where single edited cells are sequenced and indel calling is limited to sites with to five mismatches with the protospacer. **b**, Number of indel calls per cell. Each control or experimental cell type underwent indel calling where the target region had up to five base mismatches with either the VEGFA or EMX1 protospacer sequences. The gRNA or control listed in the key specify which gRNA that cell received. Instances where the indel is called in a genomic region that does not match the gRNA received by that cell are presumed to be false positives. **c**, Table of total number of off-target indel locations called that were either unique to one cell or found in multiple cells **d**, Genomic locations of recurrent indels with EMX1 or VEGFA gRNAs. On-target sites are noted in gray. **e**, Circos plots of SV identified in each cell type that received either the EMX1 or VEGFA gRNA with sites that contained at least one recurrent breakpoint seen across cell types in green or only in that cell type in red. The number of SV detected per cell are plotted to the right. (for boxplots center line is the median; box limits represent upper and lower quartiles; whiskers represent 1.5x interquartile range; points show outliers)

Compared to control cells that either received Cas9 alone or had a mock transfection, as expected there were more off-target indels in the VEGFA edited cells that showed wide cell-to-cell variance while only a small number of off-target EMX1 editing events were detected (Fig. 6b). It was noted that most of the presumed false positive edits that were seen in the control cells were single base pair insertions. Removal of non-recurrent single base pair insertions further improved the precision of indel calling (Supplementary Fig. S15). To further validate our off-target indel calling strategy, we also performed targeted deep sequencing of the PTA product from ES cells where we were able to validate 5 VEGFA off-target events (Supplementary Fig. S16). Interestingly, we found that most, but not all, recurrent off-target sites were cell-type-specific, further supporting the finding that the general chromatin structure of a cell type influences off-target genomic locations (Fig. 6d)(38).

We then performed structural variant (SV) calling to identify genome editing-induced SV where we again required regions around both breakpoints have a perfect match to the PAM sequence and allowed up to 5 mismatches with the protospacer. We again measured increased numbers of SV with the VEGFA guide RNA, with only one SV detected in EMX1-edited cells and no SV detected in control cells (Fig. 6e). We also detected recurrent VEGFA-mediated SV, some of which were cell-type specific. Interestingly, we detected more SV in the ES cells; additional studies are needed to determine if ES cells are more prone to CRISPR-mediated SV.

## Discussion

Here we present PTA, a new WGA method that captures a larger percentage of the genomes of single human cells in a more uniform, accurate, and reproducible manner than previous approaches. This results in a greatly improved capacity to SNV, as well as CNV and other SV of genetic variation in each cell, allowing for new types of studies that measure genetic variation within a tissue or organism at single-cell resolution. Importantly, PTA has far superior SNV detection sensitivity than all existing methods which is important for applications such as preimplantation genetic testing and cancer cell genome profiling where the frequent random loss of critical biological variants would significantly hamper the ability to perform those studies. In addition, we show that we can also focus on the highest quality PTA variant calls to estimate per cell mutation rates by extrapolating the total number of somatic variants across the genome based on the percentage of germline variants detected. This has important advantages over single-cell colony sequencing that could select for certain cells and is impractical for many types of primary cells. Still, future studies could make further improvements to the variant calling tools, including that incorporation PTA-specific artifacts for SNV, Indel, and SV calling, as well as including the detection of other genomic features such as mobile element insertions. In addition, PTA could be combined with whole transcriptome amplification and protein abundance in the same cells to deconvolute the contributions of cell state and genotype to a given cellular phenotype(39).

To demonstrate the potential utility of PTA for new research applications, we developed DMEM, a human mutagenicity assay that provides a framework for studying interactions between distinct environmental exposures and a living human genome. With this assay, we were able to measure the magnitude, genomic locations, and signature of ENU-induced mutations in living human cells. In future studies, it will be important to determine if mutagen signatures are cell-type specific, as well as any changes that may occur to the signature when a given mutagen is altered by metabolism by normal cells, including hepatocytes. Still, this provides an opportunity to begin to catalogue the signatures of mutagens at much higher resolution in living human cells.

We also developed an approach for measuring the off-target activity of genome editing strategies at single-cell resolution. We found that recurrent off-target editing that produces indels or SVs can be cell-type-specific, suggesting that evaluations of off-target gRNA activity should be done in the context of a specific cell type of interest. These findings also establish the general feasibility for performing single-cell biopsies of edited tissues to determine the fidelity and safety of a given genome editing intervention. Still, additional work is needed to improve the variant calling tools, as indels are difficult to call in general, and chimeras produced by WGA methods can give false positive SV calls.

DMEM and the measurement of off-target genome editing activity are just the first of a number of applications that are now possible with PTA. Other examples include more accurate preimplantation genetic testing to limit the inheritance of life-threatening genetic diseases(40), estimating cancer population diversity through the accurate measurements of the per cell mutation burdens, determining cancer-therapy induced mutagenesis rates(41), and further establishing the contributions of somatic mosaicism to human health(42). In addition to estimating genetic diversity in multicellular organisms, the improved performance of PTA could also produce more accurate and complete genome assemblies of novel unicellular organisms(43). Finally, PTA is amenable to high-throughput, parallelized reactions in microfluidic devices or emulsions. Established microfluidic strategies could be adpopted for high-throughput PTA, such as those used for digital droplet MDA (44, 45). PTA provides the technological framework for studying the genomic diversity and evolution of individual eukaryotic cells, which will undoubtedly provide new insights into tissue health and disease.

## Methods

### PTA Development Experiments

Lymphoblastoid cells from 1000 Genomes Project subject GM12878 (Coriell Institute, Camden, NJ, USA) were maintained in RPMI media, which was supplemented with 15% FBS, 2 mM L-glutamine, 100 units/mL of penicillin, 100 μg/mL of streptomycin, and 0.25 μg/mL of Amphotericin B (Thermo Fisher). The cells were seeded at a density of 3.5 × 10^5^ cells/ml and split every 3 days. They were maintained in a humidified incubator at 37°C with 5% CO_2_. Prior to single cell isolation, 3 mL of suspension of cells that had expanded over the previous 3 days was spun at 300xg for 10 minutes. The pelleted cells were washed three times with 1 mL of cell wash buffer (1X PBS containing 2%FBS without Mg2 or Ca2) where they were spun sequentially at 300xg, 200xg, and finally 100xg for 5 minutes to remove dead cells. The cells were then resuspended in 500 *μ*L of cell wash buffer, which was followed by viability staining per the manufacturer’s instructions with 100 nM of Calcein AM (Molecular Probes) and 100 ng/ml of propidium iodide (Sigma-Aldrich). The cells were loaded on a BD FACScan flow cytometer (FACSAria II) (BD Biosciences) that had been thoroughly cleaned with ELIMINase (Decon Labs,) and calibrated using Accudrop fluorescent beads (BD Biosciences). A single cell from the Calcein AM-positive, PI-negative fraction was sorted in each well of a 96 well plate containing 3 μL of PBS (Qiagen, REPLI-g SC Kit) with 0.2% Tween 20. Multiple wells were intentionally left empty to be used as no template controls. Immediately after sorting, the plates were briefly vortexed and centrifuged, and immediately placed on dry ice. Cells were then stored at −80°C for a minimum of 8 hours until ready to use.

### PTA and SCMDA Experiments

WGA Reactions were assembled in a pre-PCR workstation that provides constant positive pressure with HEPA filtered air and which was decontaminated with UV light for 30 minutes before each experiment. MDA was carried according to the SCMDA methodology using the REPLI-g Single Cell Kit (Qiagen) according the published protocol(16). PTA was carried out by first further lysing the cells after freeze thawing by adding 2 *μ*L a prechilled solution of a 1:1 mixture of 5% Triton X-100 (Sigma-Aldrich) and 20 mg/ml Proteinase K (Promega). The cells were then vortexed and briefly centrifuged before placing at 40°C for 10 minutes. 4 *μ*L of Buffer D2 (REPLI-g Single Cell Kit, Qiagen) and 1 *μ*L of 500 *μ*M exonuclease-resistant random primer were then added to the lysed cells to denature the DNA prior to vortexing, spinning, and placing at 65°C for 15 minutes. 4 *μ*L of room temperature Stop solution (REPLI-g Single Cell Kit, Qiagen) was then added and the samples were vortexed and spun down. 56 *μ*L of amplification mix (REPLI-g Single Cell Kit, Qiagen) that contained alpha-thio-ddNTPs (Trilink Bio Technologies) at equal ratios at a concentration of 1200 *μ*M in the final amplification reaction. The samples were then placed at 30°C for 8 hours after which the amplification was terminated by heating to 65°C for 3 minutes. After the SCMDA or PTA amplification, the DNA was purified using AMPure XP magnetic beads (Beckman Coulter) at a 2:1 ratio of beads to sample volume and the yields were measured using the Qubit dsDNA HS Assay Kit with a Qubit 3.0 fluorometer according to the manufacturer’s instructions (Thermo Fisher).

### Library Preparation

1ug of SCMDA product was fragmented for 30 minutes according to the KAPA HyperPlus protocol (Roche). The samples then underwent standard library preparation with 15 *μ*M of unique dual index adapters (Integrated DNA Technologies) and 4 cycles of PCR. The entire product of each PTA reaction was used for DNA sequencing library preparation using the KAPA HyperPlus kit without fragmentation. 2.5 *μ*M of unique dual index adapter (Integrated DNA Technologies) was used in the ligation, and 15 cycles of PCR were used in the final amplification. The libraries from SCMDA and PTA were then visualized on a 1% Agarose E-Gel (Invitrogen). Fragments between 400-700 bp were excised from the gel and recovered using the ZymoClean Gel DNA Recovery Kit (Zymo Research). The final libraries were quantified using the Qubit dsDNA BR Kit (Thermo Fisher) and Agilent 2100 Bioanalyzer (Agilent Technologies) before sequencing on the NovaSeq 6000 (Illumina).

#### Primary Cell Commercial WGA Kit Comparison

To compare PTA to other WGA kits that have also recently become available commercially, we performed PTA per the manufacturer’s recommendations (BioSkryb Genomics), which includes performing all initial lysis and neutralization steps on ice. Other specific recommendations for cell sorting and reproducible genome amplification can be found on the website (https://www.bioskryb.com/resources/). The five other commercially available single cell WGA kits used in the comparison were: Ampli1 (Menarini Silicon Biosystems), PicoPlex Gold (Takara Bio), Single Cell GenomiPhi (GE Healthcare), TruePrime (Expedeon), and REPLIg Single Cell (Qiagen). The study was performed on five primary leukemia samples with known copy number variation based on bulk sequencing. The deidentified leukemia samples were obtained from the St. Jude tissue bank using the an IRB-approved protocol that included the consent of the patient or patient’s parents in accordance with the Declaration of Helsinki. For this study, we first thawed and sorted a single vial of a primary childhood acute lymphoblastic leukemia sample using the procedures outlined above. Forty plates of single cells were frozen at −80 until they were ready for use. In each plate, one of the WGA protocols were assembled using a dedicated DNA-free pre-PCR laminar-flow hood. All reactions were carried out according to manufacturer’s instructions with the kit-specific changes noted below. Unless otherwise specified, 500ng of input were used for library preparation with 30 minutes of fragmentation and 8 PCR cycles, and the products were size selected using a double-sided AMPure bead (Beckman Coulter) purification (0.55X/0.25X). Final library fragment size distributions and concentrations were determined using the Qubit High Sensitivity Assay (Thermo Fisher) with the Bioanalyzer 2100 (Agilent) or the D1000 ScreenTape Assay with the TapeStation 2200 (Agilent).

We performed the following modifications for specific kits: Ampli1 reaction volumes were increased proportionately to match the 3ul starting volume and were then followed as per the manufacture’s instructions. As per the manufacturer’s recommendation, to increase the total dsDNA content, we used the Ampli1 ReAmp/ds kit and removed the adaptors with MseI (New England Biolabs). As per their recommendation, we also used the Ampli1 QC kit to select products that were positive for four PCR markers, SMARTer-PicoPlexGold reaction volumes were also increased proportionaley to accommodate the 3ul starting volumes, and the remaining steps, including library preparation, were followed as per manufacturer’s instructions using the SMARTeR Unique Dual Indexing Kit (Takara).

### Benchmarking Experiments Data Analysis

Data were trimmed using trimmomatic(46) to remove adapter sequences and low-quality terminal bases, which was followed by GATK 4.1 best practices with genome assembly GRCh38. All files were downsampled to the specified number of reads using Picard DownSampleSam. Quality metrics were acquired from the final bam file using qualimap, as well as Picard AlignmentMetricsAummary and CollectWgsMetrics. Lorenz curves were drawn and Gini Indices calculated using htSeqTools in R. SNV calling was performed using HaplotypeCaller, and subsequent VCF files were filtered using Tranche 99.0. No regions were excluded from the analyses and no other data normalization or manipulations were performed.

#### Coefficient of Variation Calculations and Copy Number Variation Calling

We first split all chromosomes into windows of specified sizes. To calculate the coverage within each window, we filtered reads to make sure MAPQ >= 40, and masked the regions with gaps (https://gist.github.com/leipzig/6123703) or segmental duplications (http://humanparalogy.gs.washington.edu/build37/data/GRCh37GenomicSuperDup.tab). We then summed up the read counts within each window to generate the maps. The CV of coverage vector *c* for a specific window size across all chromosomes was then calculated using

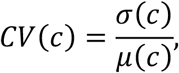

where *σ*(*c*) and *μ*(*c*) are the estimated standard deviation and mean of the coverage vector *c*, respectively.

MAPD was calculated using: 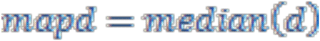, where 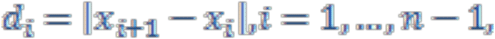, and 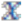 is the log2 transformed coverage of 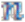 consecutive regions. If there are gaps between regions, the vector 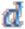 is calculated on each set of consecutive regions and then pooled together.

Ginkgo was used for CNV calling(47) after subsampling files to 10 million single reads using Picard DownSampleSam with a bin size of 1 Mbp and independent segmentation.

#### CB CD34+ Cell Enrichment for Kindred Cell and DMEM Experiments

Individual fresh anticoagulated human umbilical CB units were obtained from the St. Louis Cord Blood Bank and processed within 24 hrs of CB collection. The CB mononuclear cell fractions were prepared by Ficoll-Paque Plus (GE Healthcare) density gradient centrifugation, and immediately enriched for CD34+ cells by using the CD34 MicroBead Kit (Milteny Biotec) as per manufacturer recommendations. Enriched CD34+ cells were counted and checked for viability using a Luna FL cell counter (Logos). Cells were resuspended in freezing media (RPMI containing 15% FBS and 10% DMSO) at a density of 0.5×10^6^ cells/mL, and stored in 1mL aliquots in liquid nitrogen vapor phase.

#### Kindred Cell Isolation

Cryopreserved CB CD34+ cells were thawed using a ThawStar system (BioCision), washed with cold PBS, resuspended in SFEM II medium containing 1X CD34+ Supplement (Stem Cell Technologies) and cultured for 24hrs in a T25 flask. Cells were then stained with a CD34 antibody (APC-R700 clone:8G12, BD Biosciences), PKH67 (Sigma), and PI (Sigma). CD34+ PKH67+ PI- cells were sorted into a 1.5mL eppendorf tube containing 250uL of CD34-expansioon media. The sorted cells were subsequently used as input for a second sort in which single cells were deposited into 384-well plate wells containing 20μl StemSpan™ SFEM II (Stem Cell Technologies) with 1X CD34+ Expansion Supplement (Stem Cell Technologies). 20ul of mineral oil (Ibidi) was added to the top of the collection wells to minimize media evaporation. The sorted single cells were cultured in a humidified tissue culture incubator at 37°C with 5%CO_2_ until the single cell clones reached the 20-30 cells (day 5).

On day five, the contents of a single well were transferred from the 384-well culture plate into a 35mm dish containing 50μl of PBS. The clone cells were then mixed well by pipetting up and down using a capillary pipette. Single cells from the expanded clone were visualized and collected using an inverted microscope and a capillary pipette. The cells were collected in ~3uL and transferred to a lobind 96 well PCR plate (Eppendorf), immediately placed on dry ice, and then stored at −80C. Kindred cells underwent PTA as per the manufacturer’s instructions (startup) with the library input amount increased to 500ng and the number of PCR cycles decreased to 8.

#### Kindred Cell Variant Calling

SNVs were again identified using GATK4 as described above. To limit variant calling to covered sites, only sites with at least 15 bulk and 5 single cell reads in at least 50% of cells were considered. Custom bash scripts were then used to identify sites that were present in the the bulk sample, one single cell, or multiple single cells, as well as any combination of the three.

### Direct Measurement of Environmental Mutagenicity (DMEM)

Expanded CB CD34^+^ cells were cultured in StemSpan SFEM (Stemcell Technologies) supplemented with 1X CD34^+^ Expansion Supplement (Stemcell Technologies), 100 units/mL of penicillin, and 100 ug/mL of streptomycin. The cells were exposed to ENU at concentrations of 8.54, 85.4, and 854 *μ*M, D-mannitol at 1152.8, and 11528 *μ*M, or 20 μL of ultrapure water (vehicle control) for 48 hours. Single cell suspensions from treated cells and vehicle control samples were harvested and stained for viability as described above. Single cell sorts were also carried out as described above. PTA was performed and libraries were prepared as per the manufacturer’s instructions (startup).

### Analysis of DMEM Data

Data acquired from cells in the DMEM experiments were again trimmed using Trimmomatic, aligned to GRCh38 using BWA (0.7.12), and further processed using GATK 4.0.1 best practices without deviation from the recommended parameters. Genotyping was performed using HaplotypeCaller where joint genotypes were again filtered using standard parameters using variants in -tranche 99.9. A variant was only considered to be the result of the mutagen if it was called in only one cell while not being found in sites with at least 15X coverage in the bulk sample. The signature of ENU was created and compared to known COSMIC signatures using SigProfiler(30).

To determine whether mutations were enriched in DNase I hypersensitivity (DH) sites in CD34^+^ cells, we calculated the proportion of SNVs in each sample that overlap with DH sites from 10 CD34^+^ primary cell datasets produced by the Roadmap Epigenomics Project (http://www.roadmapepigenomics.org/). DH sites were extended by 2 nucleosomes, or 340 bases, in either direction. Each DH dataset was paired with a single cell sample where we determined the proportion of the human genome in each cell with at least 10x coverage which overlapped with a DHS, which was compared to the proportion of SNVs that were found within the covered DH sites.

#### CRISPR Off-Target Editing Measurements

We edited three cell types: cell line U20S, CB CD34+ cells, and ES cell line H9. The previously published EMX1 and VEGFA-s2 gRNA sequences were used to design the gRNAs(35). U2OS cells were maintained in DMEM (Thermo Fisher), and pools of 3×10^5^ cells were transiently co-transfected with precomplexed RNPs consisting of 150 pmole of sgRNA (Synthego) and 50 pmole of spCas9 protein (St. Jude Protein Production Core) with pMaxGFP plasmid as a transfection control at 200ng/ul (Lonza) using the 4D-Nucleofector™ X-unit (Lonza) with solution P3 and program CM-104 in 20ul cuvettes according to the manufacturer’s recommended protocol. CB cells were maintained in SFEM II medias described above. For the CB RNP transfection, 1.6×10^5^ cells were suspended in P3 Primary Cell Solution with 50pmol Cas9 protein (St. Jude Protein Production Core) and 150pmol sgRNA (Synthego) with pMaxGFP plasmid at 200ng/ul (Lonza) in a total volume of 20ul with program DS130. 1×10^6^ H9 ES cells were pretreated with mTeSR1 (Stem Cell Technologies) supplemented with 1X RevitaCell (Thermo Fisher) for 2 hours, and then transiently co-transfected with 500 pmol of sgRNA (Synthego),168 pmol of spCas9 protein (St. Jude Protein Production Core), and pMaxGFP plasmid at 200ng/ul (Lonza) using solution P3 and program CA-137.

Cells were all plated and maintained in the same media for 48-72 hours prior to sorting, and matrigel (Corning) coated plates were used for recovering the ES cells. A FACS Aria flow cytometer (BD Biosciences) was used to collect PI negative, GFP positive single cells as described above. >10,000 cells were also collected in a single tube as a bulk control. Sorted single cell underwent PTA, library preparation, and sequencing as described for the kindred cell experiment. Following library preparation, targeted PCR followed by sequencing was used to confirm editing at the expected site for each gRNA.

### CRISPR Single Cell Analyses

Small indels were also identified using GATK version 4 best practices, selecting variants in tranche 90.0 to limit the number of false positive indel calls. To limit our search to covered genomic regions, we again required the bulk samples sites have at least 15X coverage to be considered, and that at least 50% of the single cells had at least 5X coverage. Indels were required to overlap regions within 50 base pairs of the canonical cleavage site of a potential off-target site for each gRNA. Potential off-target sites for each gRNA were identified using Cas-Offinder(48) to find all 23mers in GRCh38 that were different at no more than 5 bases from the 20mer protospacer of the gRNA and that matched NGG in the adjacent PAM sequence. Only potential off-targets from chr1-22, chrX, chrY, and chrM were considered, and indels that were called in the bulk sample or any of the other cells that did not receive that gRNA were also excluded from consideration. Large indels and other structural variants were identified using svABA(49), which identifies and outputs paired breakpoints. Breakpoints were identified from the same bam files that were used for small indel calling. SvABA was run separately for each treatment type in each gRNA experiment using the case-control option. For example, to identify structural variants in VEGFA-edited CD34 cells, the VEGFA-edited CD34 cells were treated as the case samples and the remaining CD34 cells were treated as controls. Only variants that were called in the case samples and were identified in at least one cell were considered further. Structural variants were only considered if both breakpoints were within 200 bases of the canonical cleavage site of a potential gRNA off-target site, allowing up to 5 mismatches while again requiring the NGG PAM sequence.

### Code Availability

The authors have used open source software to perform all analysis. To ensure our results are reproducible, we have made scripts needed to reproduce our results available on request.

---

**Supplementary Fig. S1.**
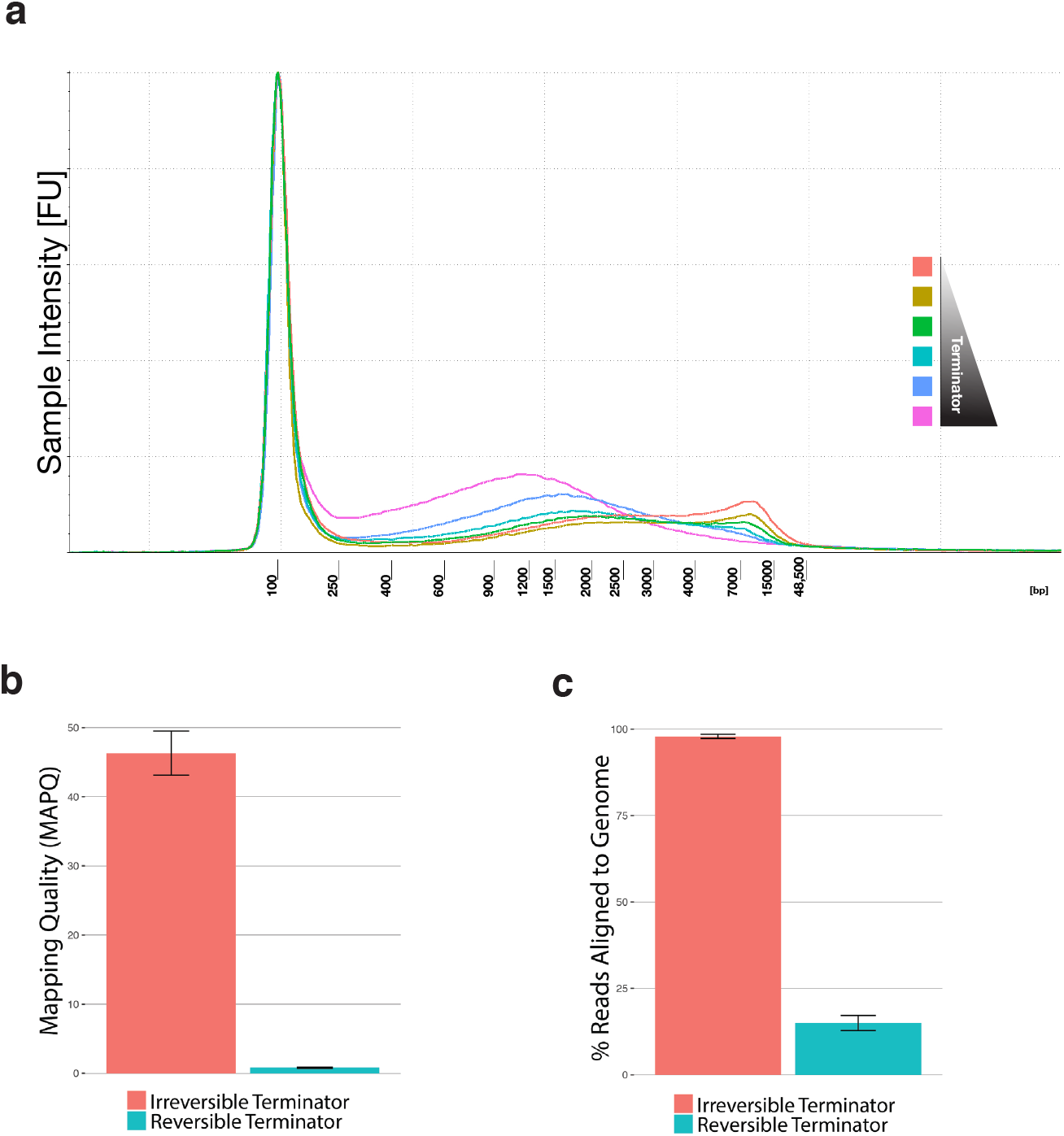
The incorporation of irreversible terminators in PTA produces smaller, high-quality WGA products. **a**, Incorporation of alpha-thio-dideoxynucleotides terminate the extension step and decrease the size of the amplification products as seen in this terminator titration experiment showing amplicon size distribution on Tapestation 4200 high molecular weight tracings. (**b** and **c**), Irreversible terminators in PTA reactions produce high-quality amplification products with a 46.3+/−3.18 mapping quality, and 97.9+/−0.62% of reads mapped to the genome, while reversible terminators produce poor quality products with an overrepresentation of repetitive elements. (error bars represent one SD)

**Supplementary Fig. S2.**
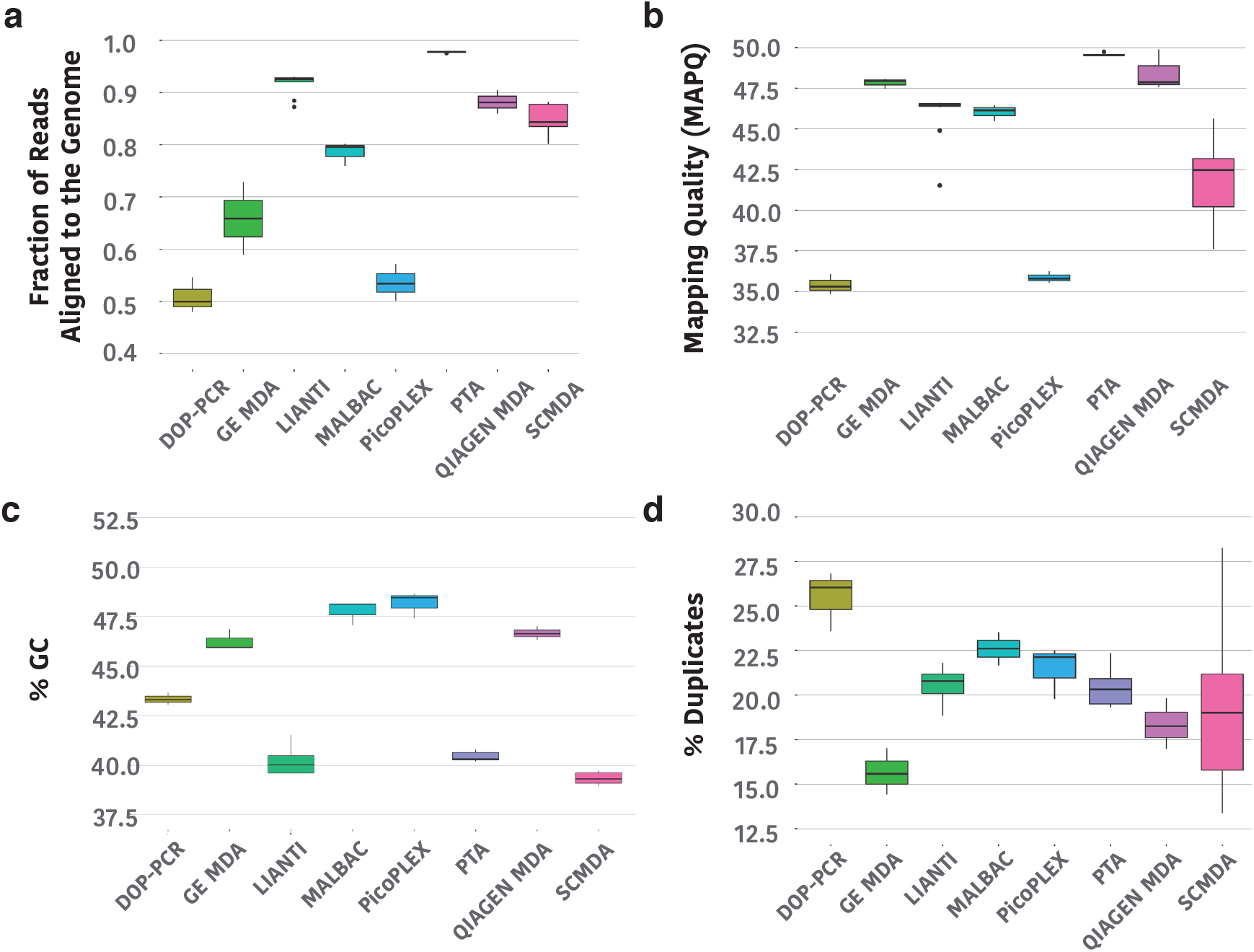
Performance comparisons of PTA to the most common single-cell WGA methods. PTA and SCMDA were performed on 10 GM12878 cells each followed by library preparation and NGS. The resulting sequencing data was compared to data produced from single cells that had undergone amplification with DOP-PCR (n=3), GE MDA (n=3), Qiagen MDA (n=3), MALBAC (n=3), LIANTI (n=11), or PicoPlex (n=3) as part of the LIANTI study. **a**, Comparison of the fraction of reads aligned to the genome. **b**, Mapping quality. **c**, Percent GC content. **d**, PCR duplication rates. PTA data has the highest percent of reads aligned to the genome, the highest mapping quality, and lower GC content than most of the other methods. PCR duplication rates were similar across all methods. (for boxplots center line is the median; box limits represent upper and lower quartiles; whiskers represent 1.5x interquartile range; points show outliers).

**Supplementary Fig. S3.**
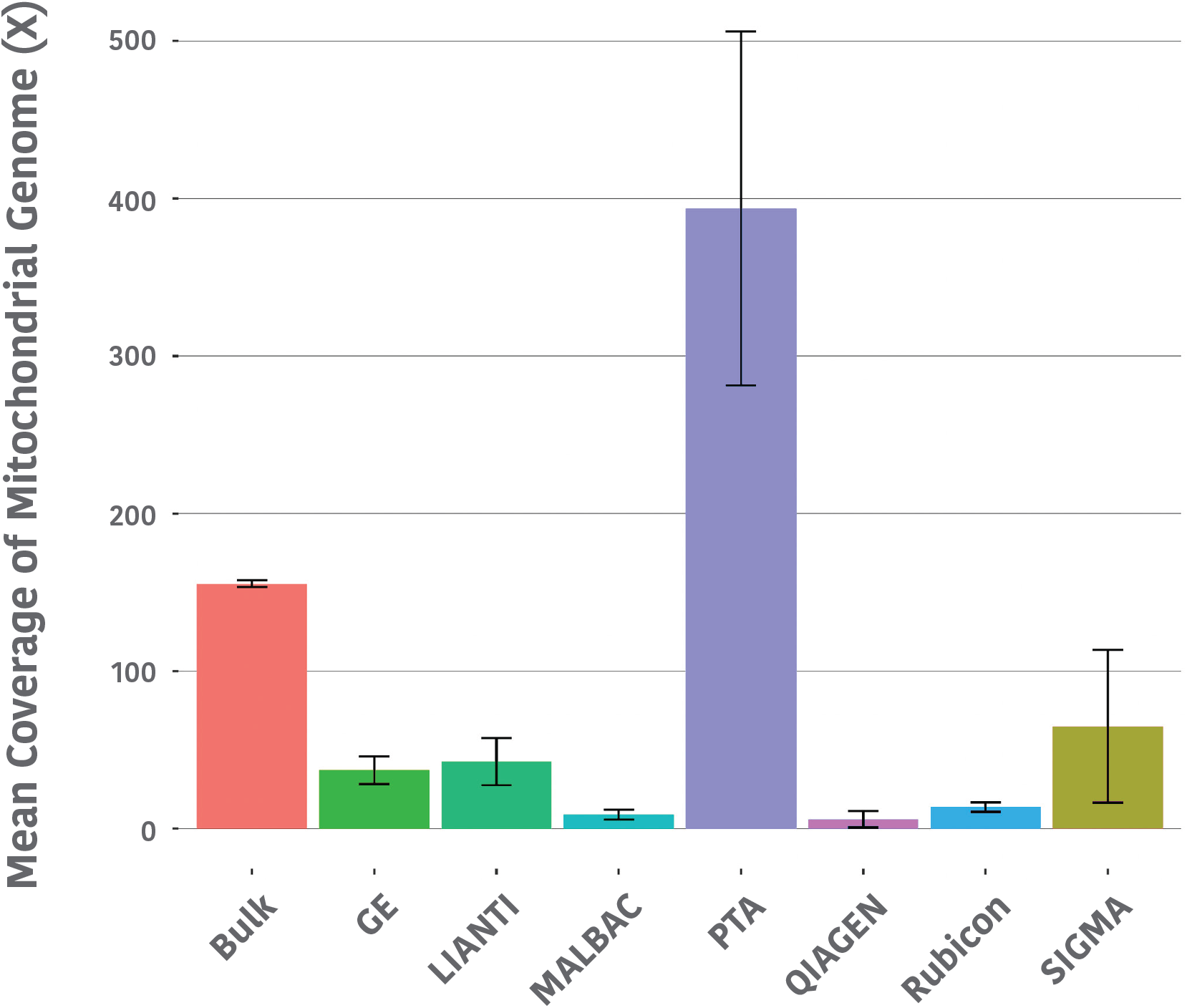
Comparison of single cell mitochondrial genome coverage breadth with different WGA methods. All comparisons were done with the same number of sequencing reads; (error bars represent one SD).

**Supplementary Fig. S4.**
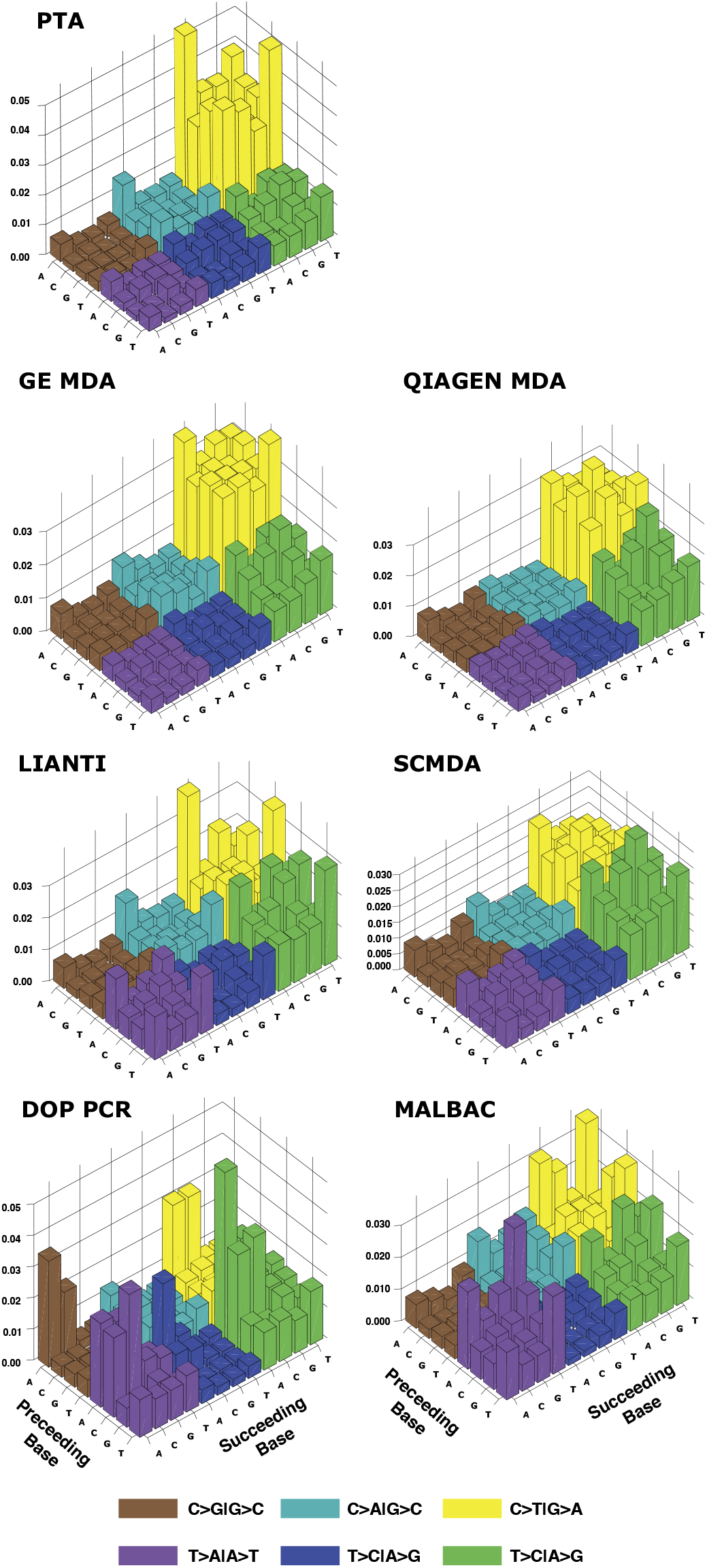
Trinucleotide base change patterns in false positive SNVs. Base change patterns seen in false positive calls appear to be polymerase-dependent with methods using an isothermal polymerase showing a preference for C to T (G to A) changes.

**Supplementary Fig. S5.**
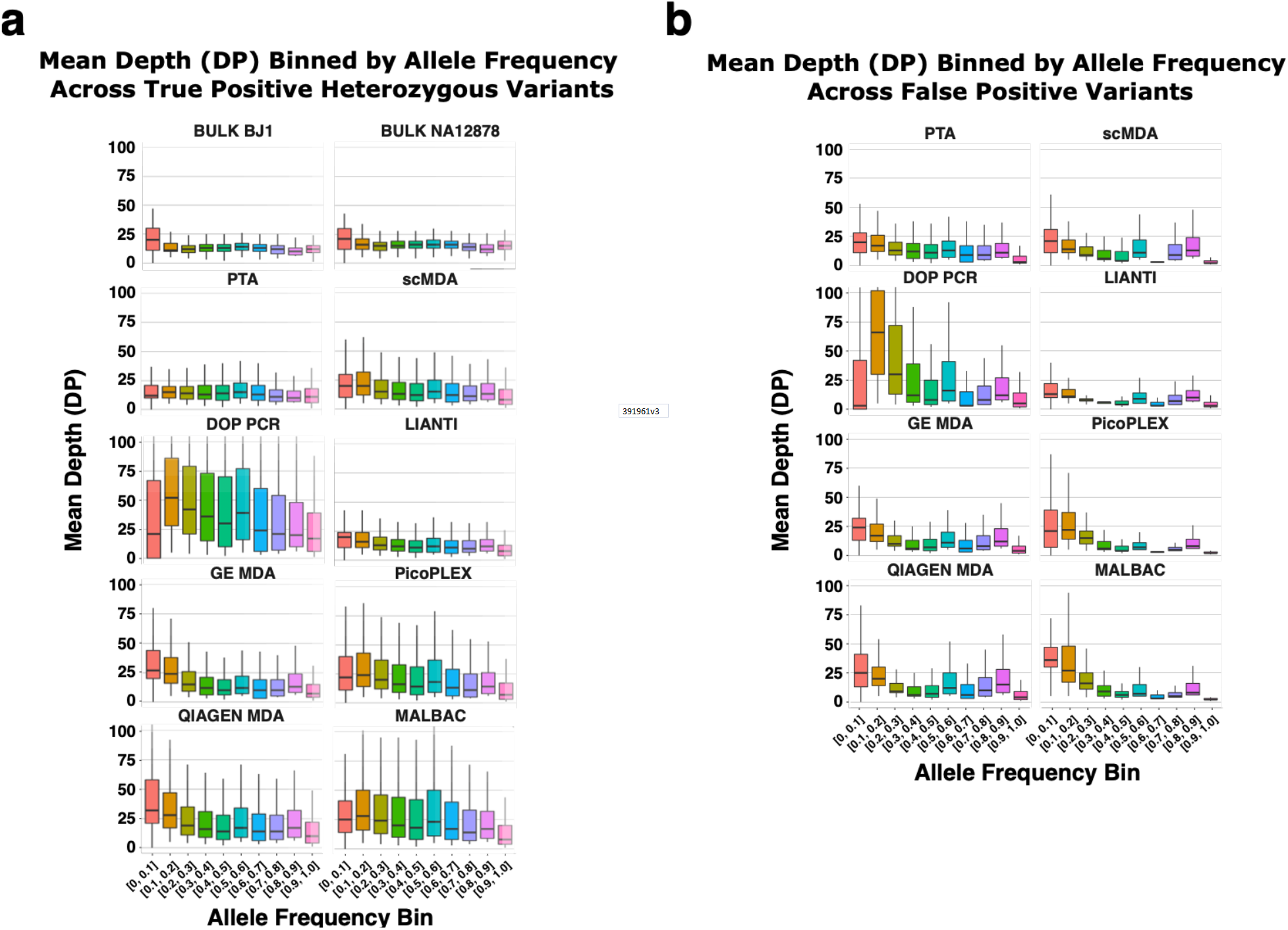
Coverage depth at increasing variant allele frequency bin for true positive (a) and false positive (b) variant calls for each of the WGA methods.

**Supplementary Fig. S6.**
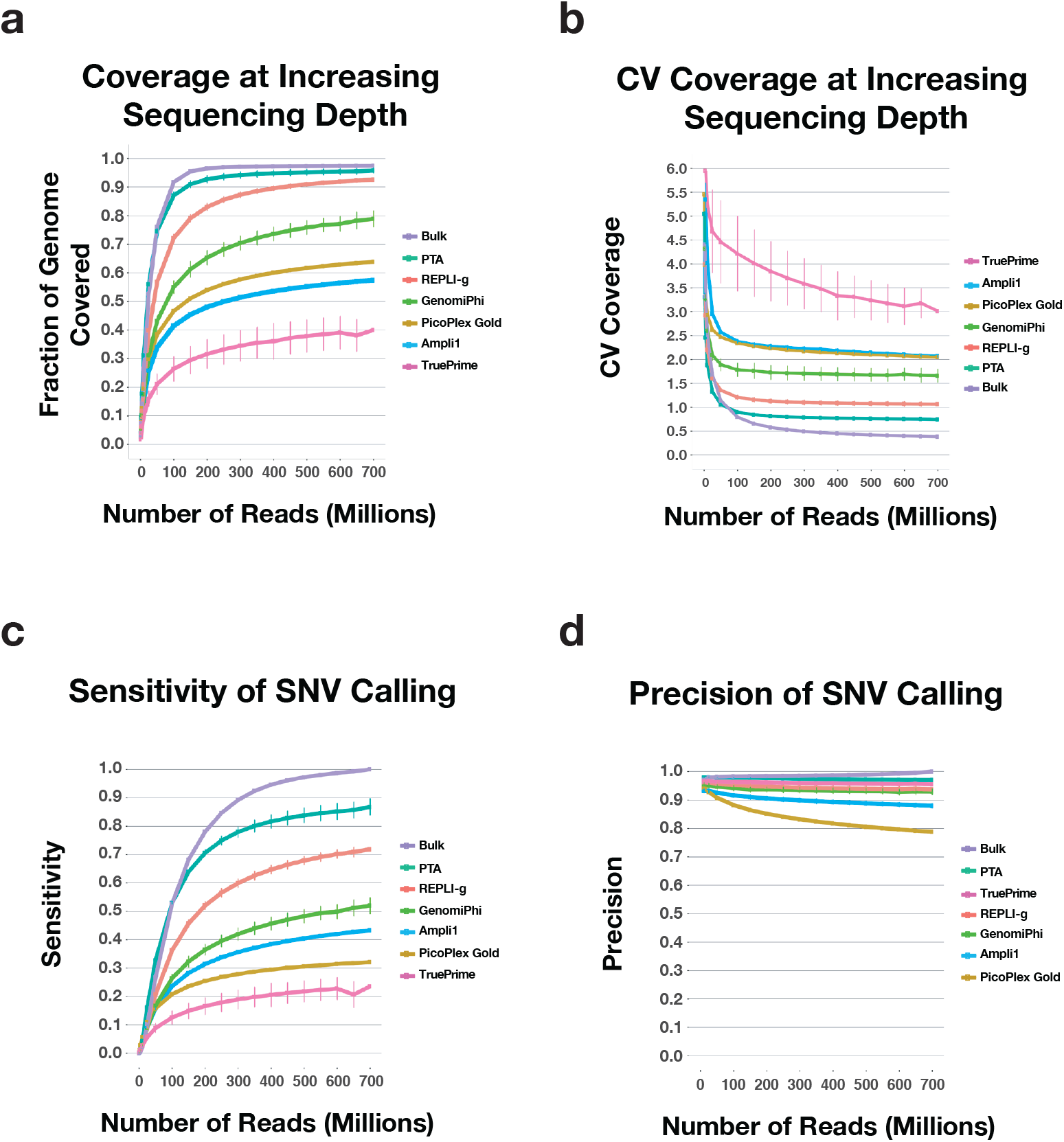
Alignment and SNV calling metrics in primary leukemia cells at increasing sequencing depth using low temperature lysis PTA. **a**, Coverage breadth **b**, CV coverage **c**, SNV calling sensitivity **d**, SNV calling precision (n=5 for each method, error bars represent one SD)

**Supplementary Fig. S7.**
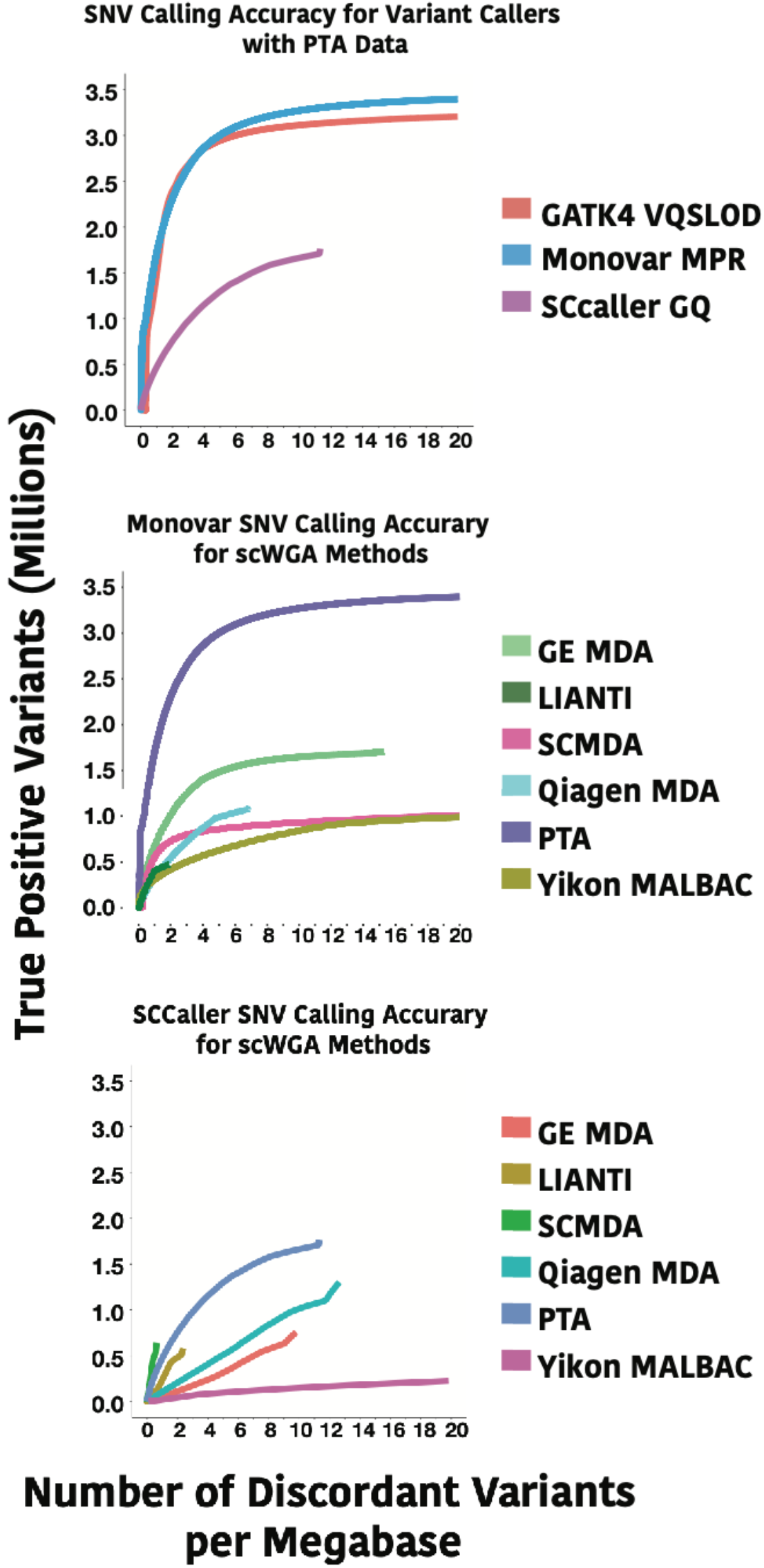
Comparison of SNV calling accuracy for different single cell variant callers.

**Supplementary Fig. S8.**
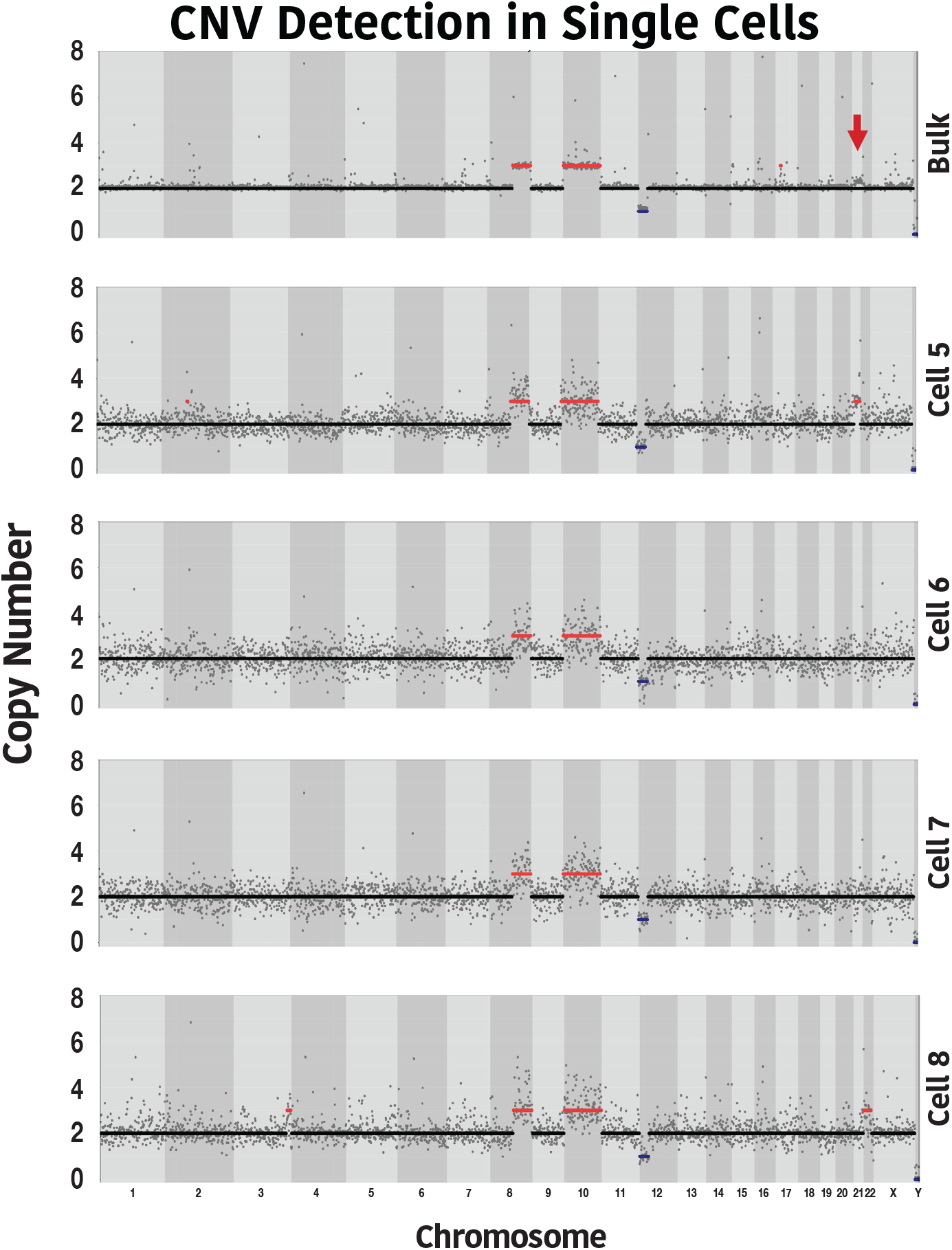
Additional single cell CNV profiles for cells presented in Fig. 3h.

**Supplementary Fig. S9.**
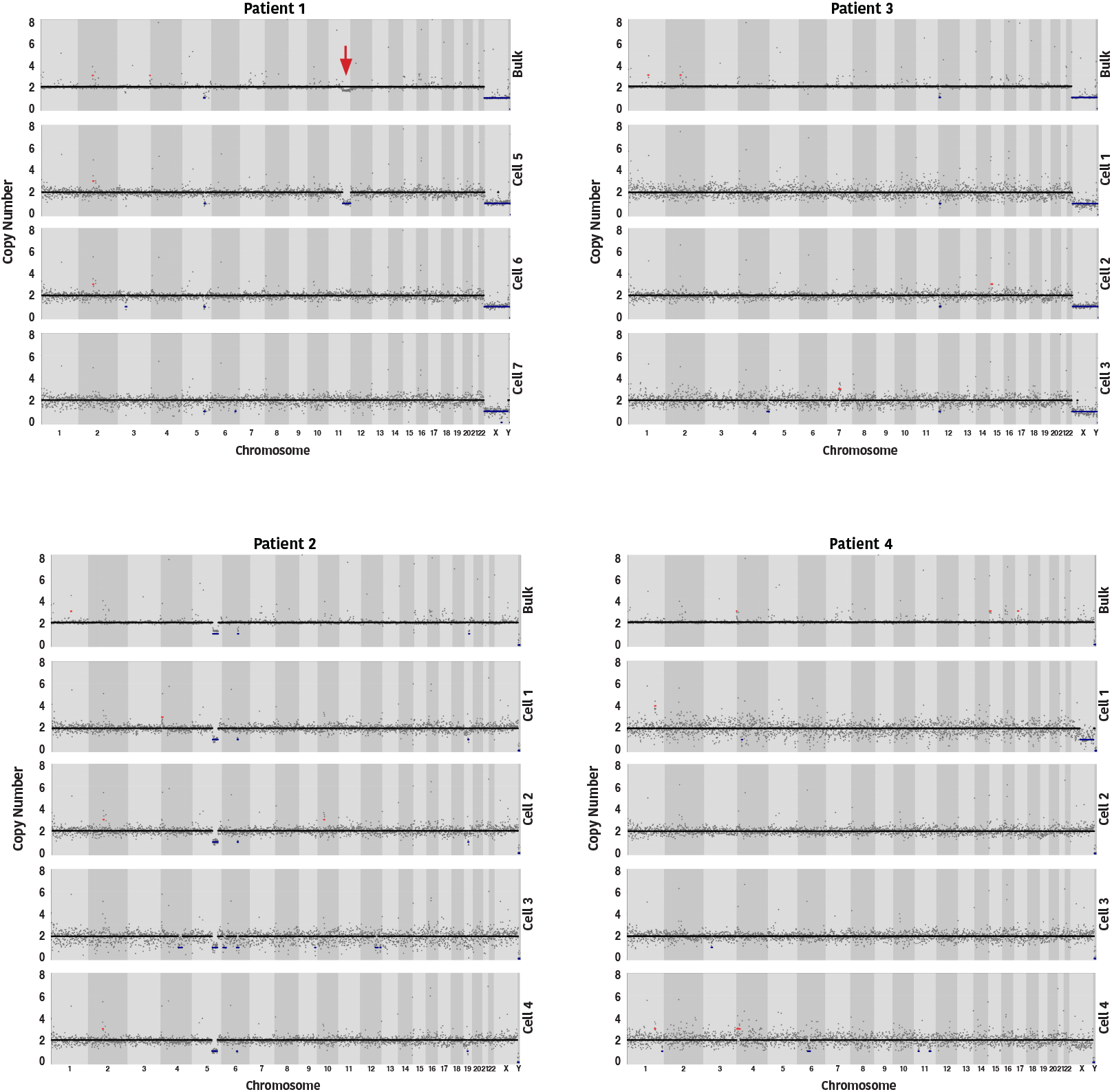
Bulk and single cell CNV profiles for four additional patient samples.

**Supplementary Fig. S10.**
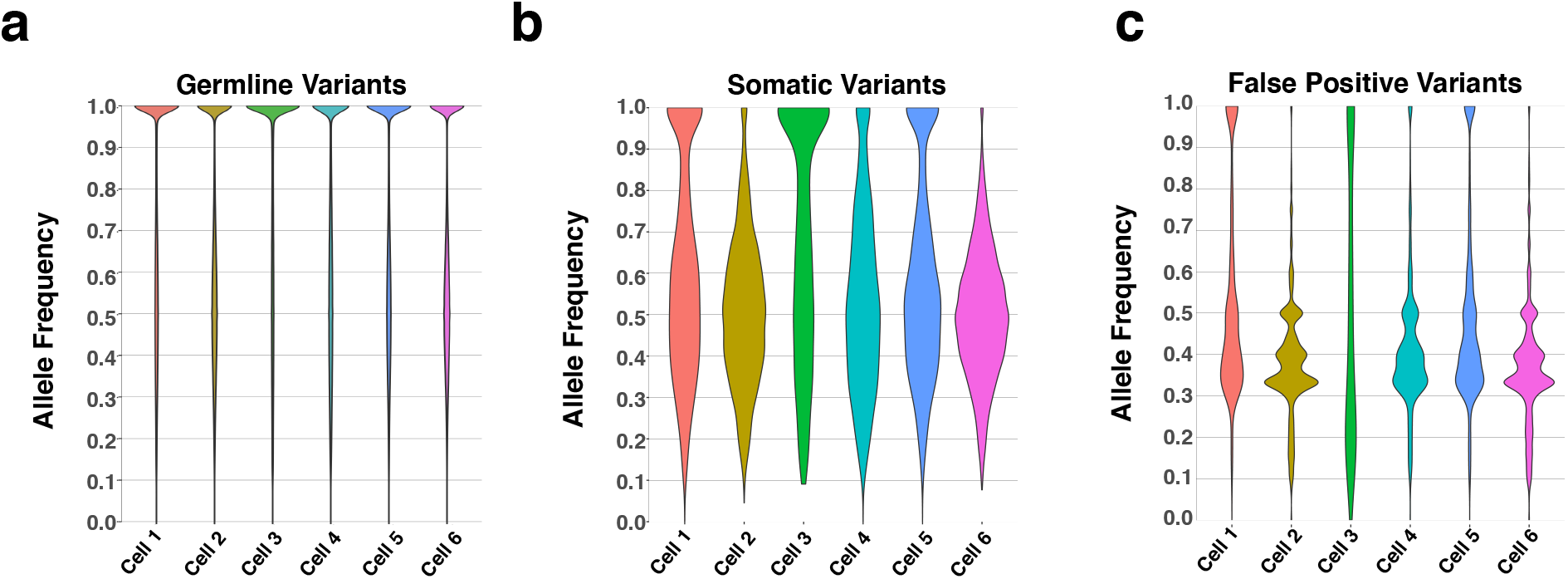
**Allele frequency distributions using standard filtering for a,** Germline variants, **b**, Somatic variants, and **c**, False positive variants.

**Supplementary Fig. S11.**
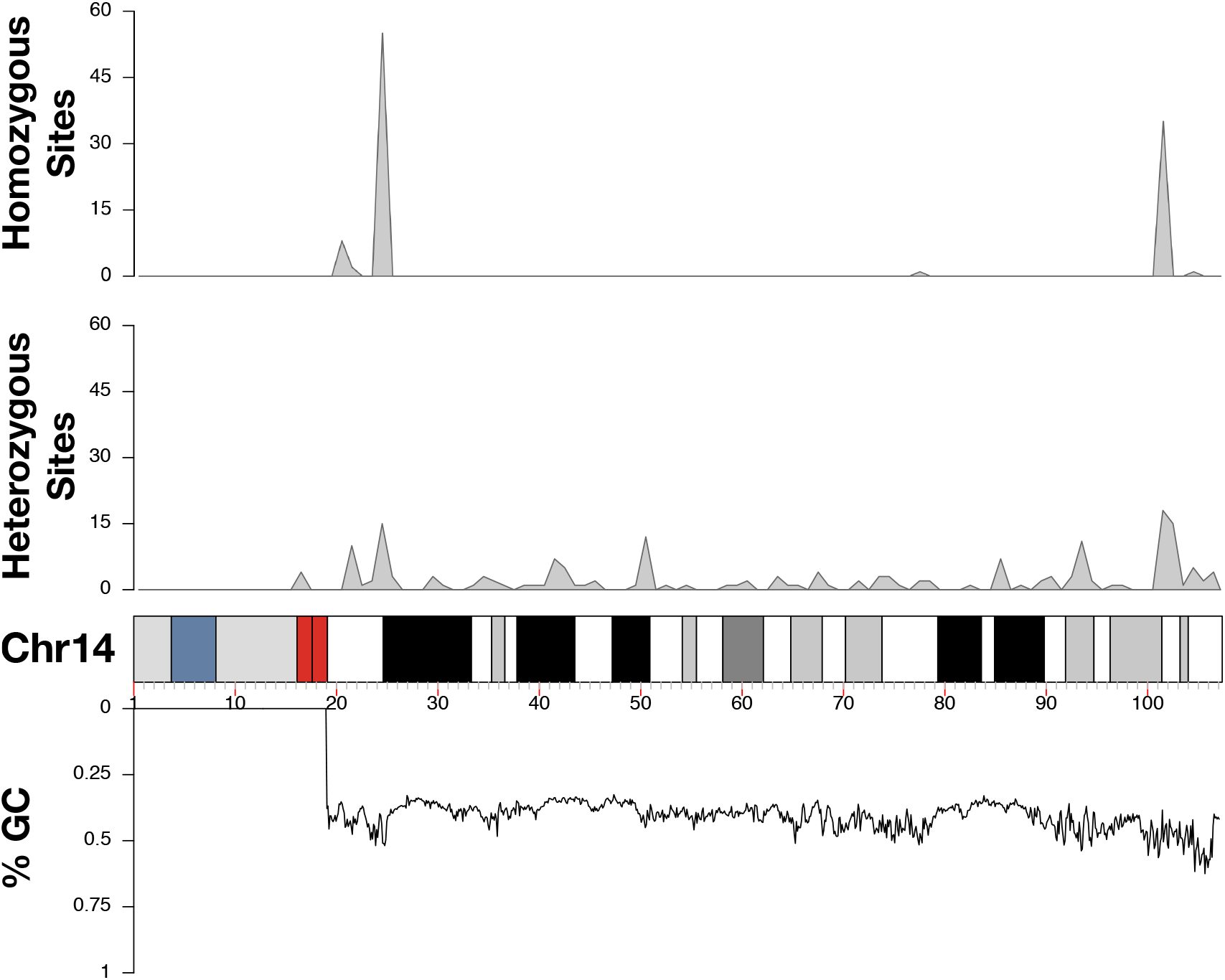
Density of homozygous or heterozygous false positive variant calls across chromosome 14. (which had the largest number of false positive calls). Mean GC content at 100 Kb intervals runs below the karyogram.

**Supplementary Fig. S13.**
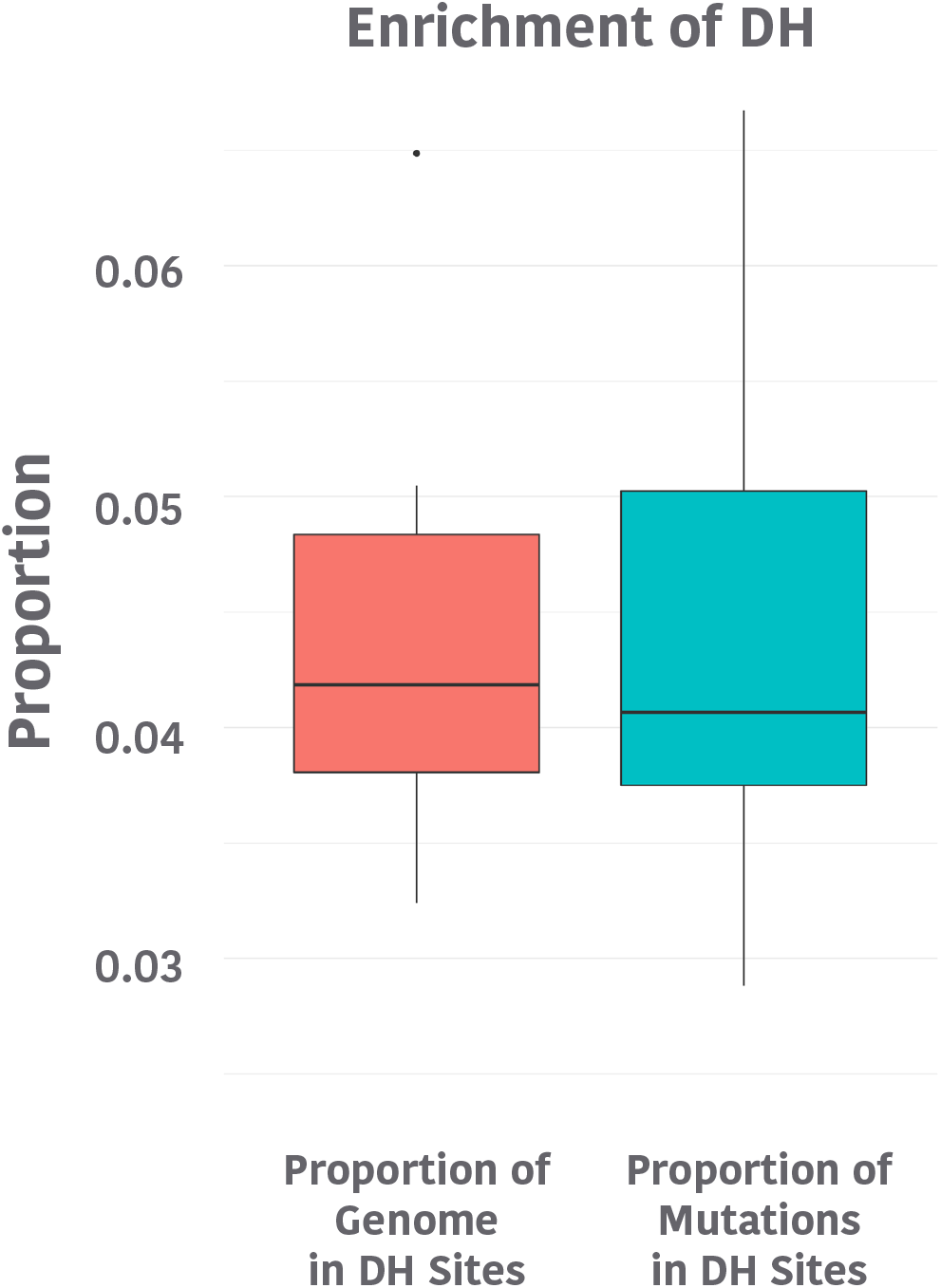
Proportion of ENU induced mutations in DNase I hypersensitive (DH) sites. **DH** sites in CD34^+^ cells previously catalogued by the Roadmap Epigenomics Project were used to investigate whether ENU mutations are more prevalent in DH sites which represent sites of open chromatin. No significant enrichment in variant locations at DH sites was identified. Further, no enrichment of variants restricted to cytosines was observed in DH sites. (for boxplots center line is the median; box limits represent upper and lower quartiles; whiskers represent 1.5x interquartile range; points show outliers)

**Supplementary Fig. S14.**
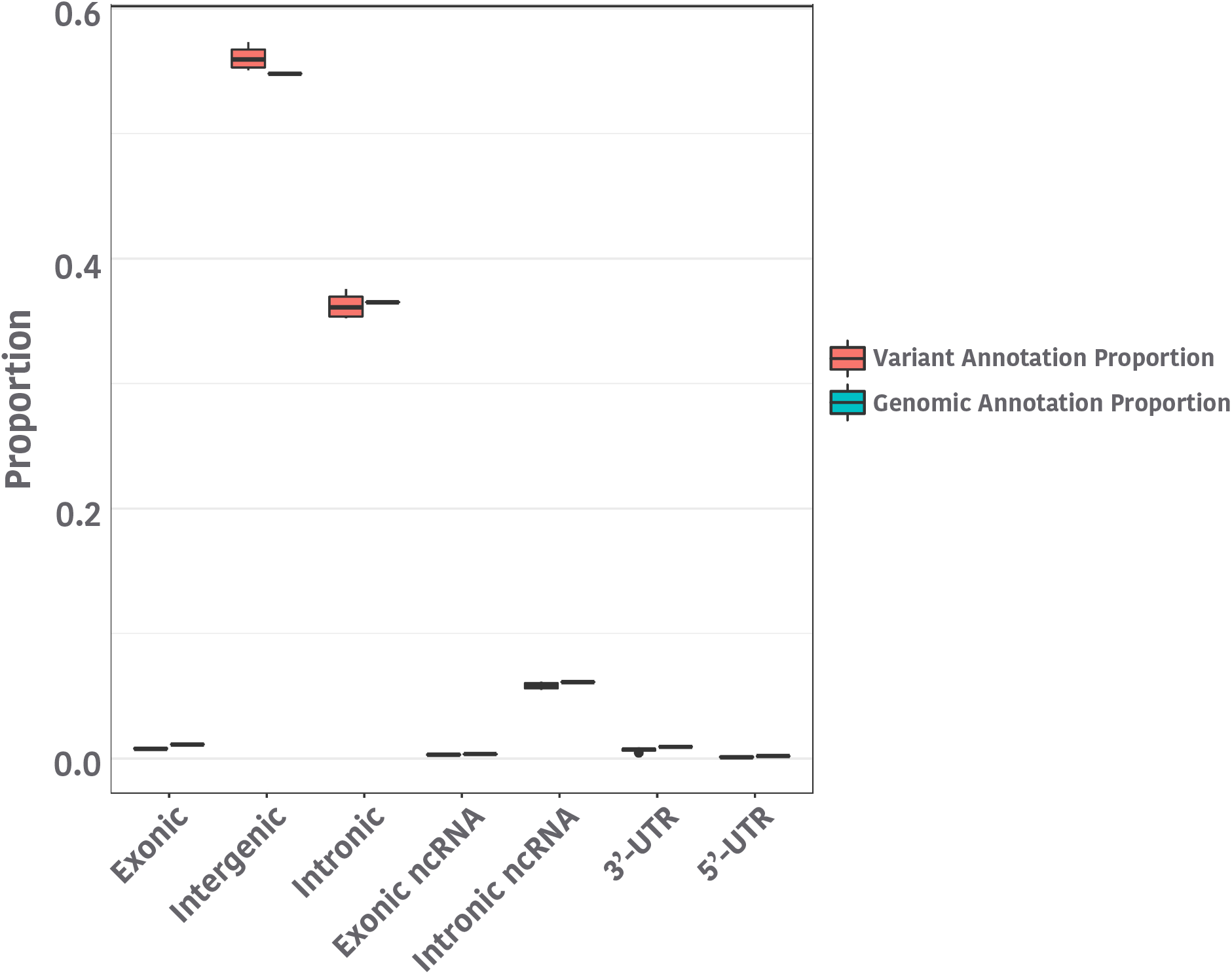
Proportion of ENU-induced mutations in genomic locations with specific annotations. No specific enrichment was seen in specific annotations for variants in each cell relative to the proportion of the genome each annotation comprises. (for boxplots center line is the median; box limits represent upper and lower quartiles; whiskers represent 1.5x interquartile range; points show outliers)

**Supplementary Fig. S14.**
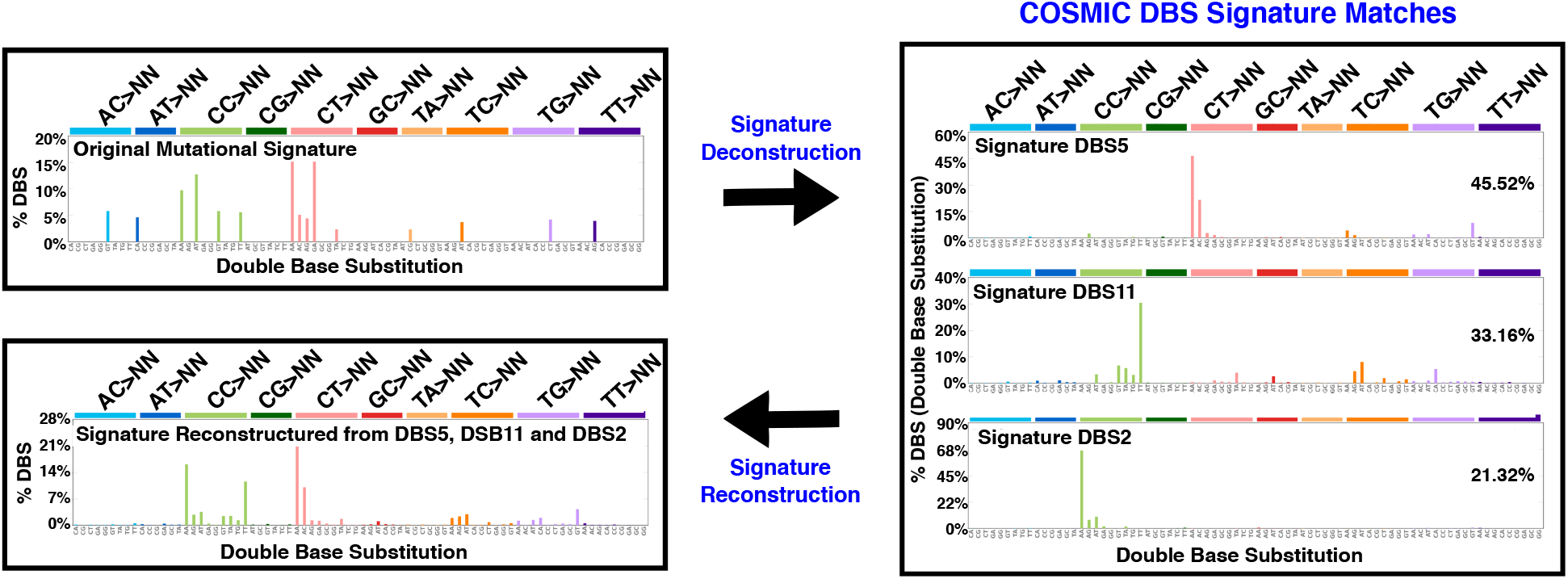
Double base substitution signature for ENU-induced variants in CD34+ cord blood cells.

**Supplementary Fig. S15.**
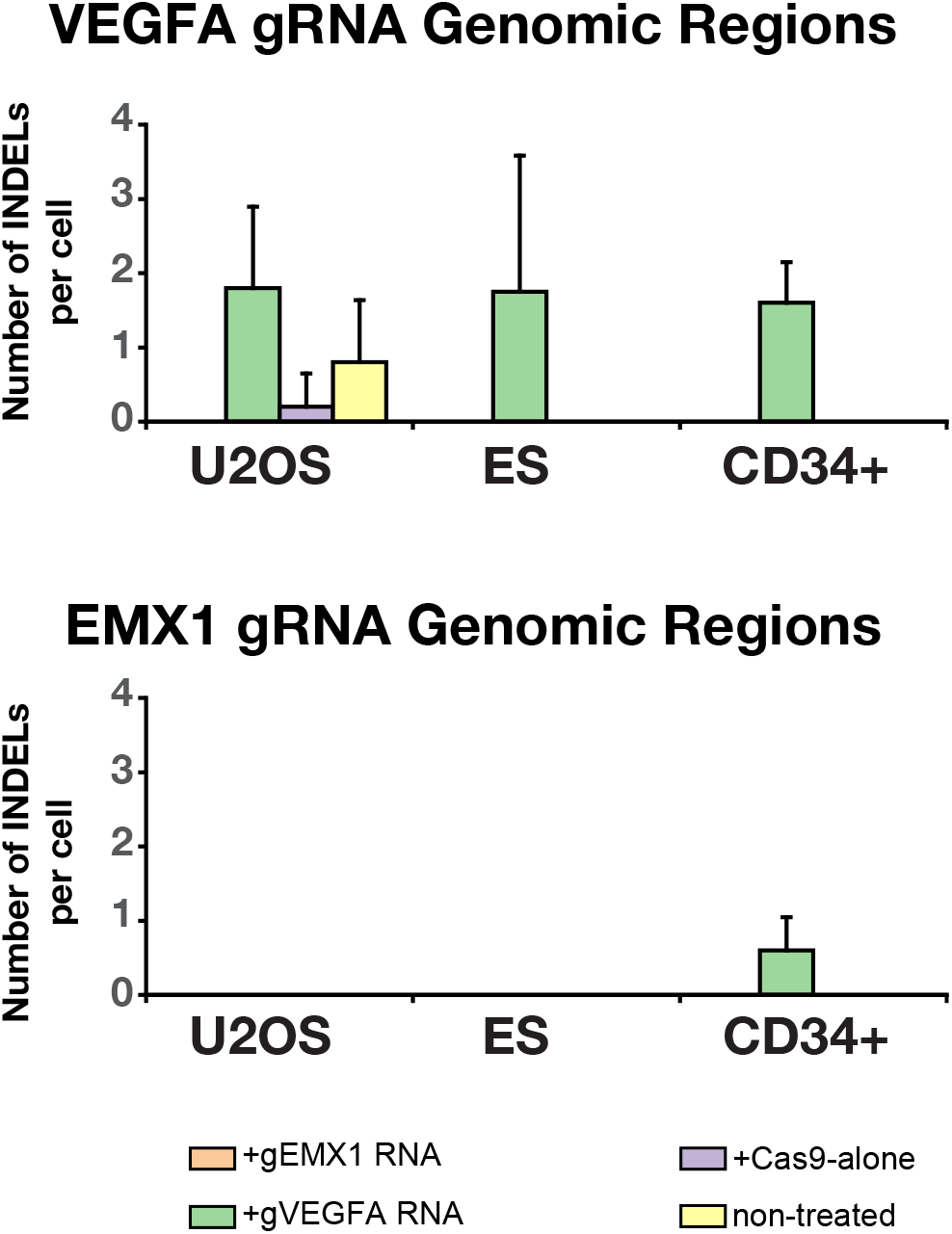
Removing non-recurrent single base pair insertions improves the precision of off-target detection. Each control or experimental cell type underwent indel calling requiring no more than five mismatches to either the VEGFA or EMX1 guide RNA sequence. Off-target events specifies which genomic region the gRNA had to match while the gRNA or control listed in the key specify which gRNA that cell received. Instances where the indel is called in a genomic region that does not match the gRNA received by that cell are presumed to be false positives.

**Supplementary Fig. S16.**
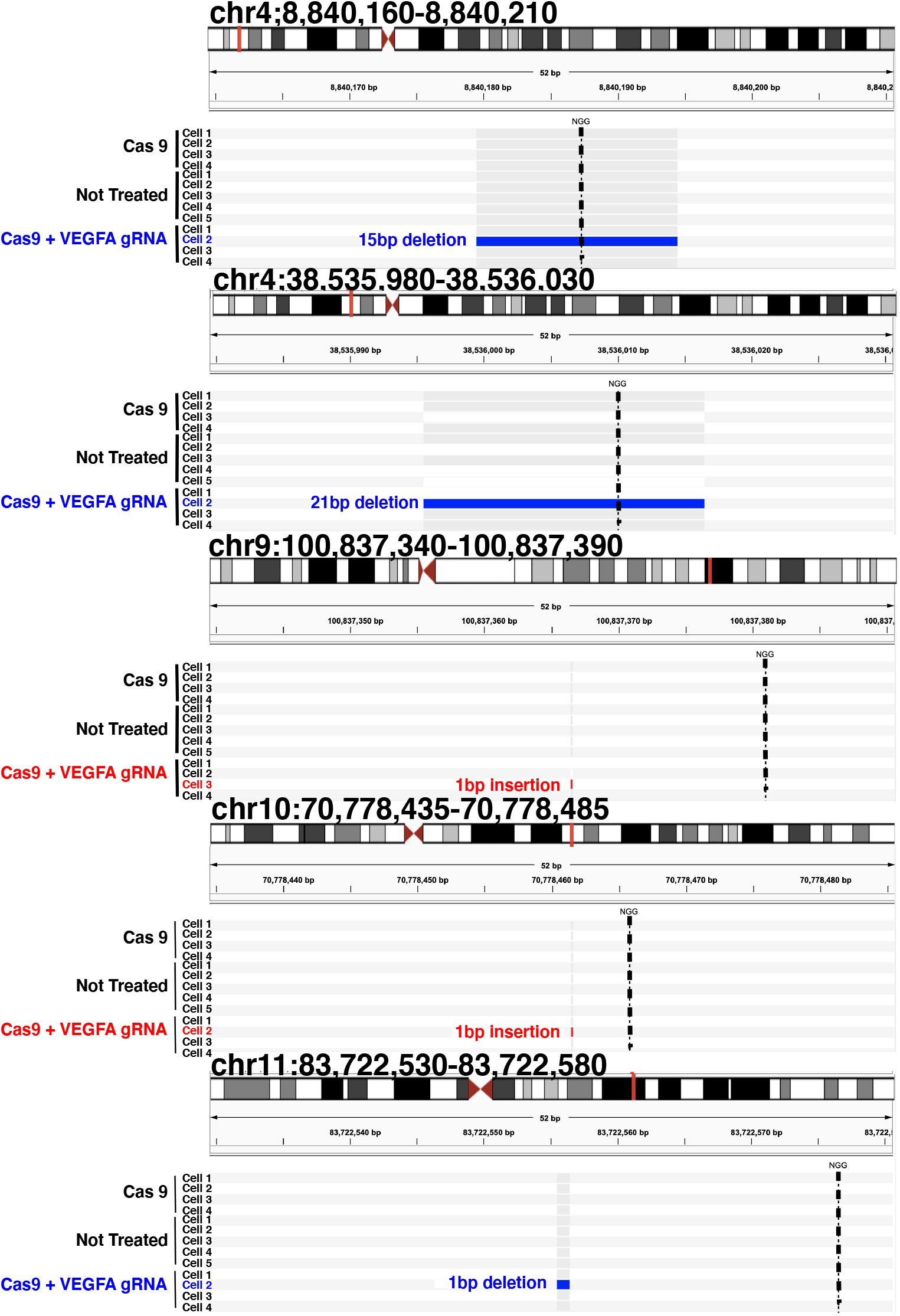
Confirmation of five VEGFA CRISPR off-target sites in ES cells using targeted resequencing.

## Acknowledgements

We would like to thank Kristine Witt and Stephanie Smith-Roe from the National Institute of Environmental Health Safety for their guidance on the design of the DMEM experiments. C.G. is supported by a Scholar Award from the Hyundai Pediatric Cancer Foundation, a Burroughs Wellcome Fund Career Award for Medical Scientists, and an NIH Director’s New Innovator Award (1DP2CA239145). We would also like to thank Josh Stokes (Biomedical Communications, St. Jude Children’s Research Hospital) for assistance with the artwork.

## Author Contributions

Conceptualization, VG, JE, CG

Methodology, VG, JE, CG

Investigation, VG, SN, KA, YP, BS, JC

Formal Analysis, TX, DK, RC, XC, DP, WC, SPM, CG

Writing– Original Draft, VG, CG

Writing– Review & Editing, VG, JE, CG

Visualization, RC, XC, YX, DK, CG

Funding Acquisition, CG

Resources, JE, CG

Supervision, JE, XC, SPM, CG

